# Abiotic Factors and Competitive Exclusion Drive Assembly Patterns in Two Insular Gecko Adaptive Radiations Displaying Ecomorphological Convergence

**DOI:** 10.64898/2026.03.02.702884

**Authors:** Phillip L. Skipwith, Nicolás Castillo-Rodríguez, Rosana Zenil-Ferguson

## Abstract

Adaptive radiation theory posits that speciation in such lineages is largely driven by ecological opportunity in concurrent morphological expansion in response to niche availability. Here, we use a phylogenomic estimate of Australasian diplodactyloid geckos in combination with meristic and ecological data to infer patterns of ecological diversification, quantify signatures of stabilizing selection, and the factors driving speciation processes. Specifically, we focus on two relatively young but speciose and ecomorphologically diverse assemblages from the ancient islands of New Caledonia and New Zealand. Models accounting for stabilizing selection recover shifts in morphospace along many branches that also experienced shifts in ecological guild as inferred from ancestral state reconstructions. We find convergent evolution to be present between the two insular lineages as they independently transitioned to similar guilds from different ancestral ecologies. Community assembly is integral to understanding the dynamics of adaptive radiations and various studies focused on identifying if biotic or abiotic factors drive character suits and sympatry in diverse lineages. Bayesian and multiple regression analyses suggest that abiotic factors rather than interspecific competition dictates phenotypic divergence in both insular lineages. Rather, species seem to diverge phenotypically in allopatry and environmental factors, such as climate, in combination with competitive exclusion drive phenotypic overlap in sympatry. This study provides the first modern assessment of convergence for diplodactyloid geckos and provides robust evidence indicating that similar selective pressures have shaped morphological diversity in these disparate as well the factors affecting sympatry.

## Introduction

Understanding patterns of taxonomic and phenotypic diversity has long been a major focus of macroevolutionary biology. Adaptive radiations (ARs), where diversity is linked to ecological opportunity, offer the possibility to contrast the tempo and mode of speciation and trait macroevolution and how these mechanisms are impacted by ecology and biogeography (Losos & Ricklefs, 2009; Schluter, 2000b; Simpson, 1944; Simpson, 1953; Wellborn & Langerhans, 2015). ARs are characterized as lineages derived from a common ancestor that speciates and diverges phenotypically in response to vacant niches. This *in situ* diversification may lead to assemblages of closely related taxa that, despite sharing common ancestral ecologies, occur in sympatry or even syntopy due to largely non-overlapping niches, minimizing interspecific competition (Gillespie, Bennett, De Meester, Feder, Fleischer, Harmon, Hendry, Knope, Mallet, Martin, Parent, Patton, Pfennig, Rubinoff, Schluter, Seehausen, Shaw, Stacy, Stervander, Stroud, Wagner & Wogan, 2020; Schluter, 2000a). In addition to this this clear ecological axis of AR, a number of authors have proposed expectations on the dynamics of speciation and trait diversification, where the two are at least partially correlated and are fastest early in the evolution of AR (Rabosky, Santini, Eastman, Smith, Sidlauskas, Chang & Alfaro, 2013; Rundell & Price, 2009). This pattern of initial rapidity with subsequent decreases in rate have been seen as a response to the reduction of viable ecological axes as niche-space contracts (Calcagno, Jarne, Loreau, Mouquet & David, 2017; Etienne & Haegeman, 2012; Etienne, Haegeman, Stadler, Aze, Pearson, Purvis & Phillimore, 2012; Pannetier, Martinez, Bunnefeld & Etienne, 2021; Rabosky, 2014; Rabosky, 2017). There is increasing emphasis on understanding the role of interspecific competition in driving character evolution, the exclusion of ecologically similar taxa from occurring in sympatry, and the importance of environmental factors in shaping geographic distributions (Gillespie et al., 2020; Pavón-Vázquez, Brennan, Skeels & Keogh, 2022; Quintero & Landis, 2019).

Biotic interactions and abiotic factors are broadly seen as having profound effects of the distributions of lineages and the evolution of niche specialization during diversification. The relative importance of abiotic and biotic drivers of diversification have been summarized in two competing hypotheses that operate on a continuum, the Red Queen (RCH) and Court Jester (CJH) hypotheses (Barnosky, 2001; Van Valen, 1973). In short, the RCH posits that diversification is driven by interactions between competing populations at relatively short timescales, while the CJH suggests that abiotic factors on longer timescales are of greater importance (Barnosky, 2001; Condamine, Rolland, Höhna, Sperling & Sanmartín, 2018). Quantifying how sympatry and ecological overlap are driven by biotic mechanisms such as character displacement and competitive exclusion versus abiotic mechanisms like environmental filtering are useful to infer how ARs are shaped.

Insular lineages are useful examples to investigate broad patterns of temporal heterogeneity in speciation and trait diversification, in addition to understanding spatial factors that may dictate niche availability and interspecific competition (Garcia-Porta, Šmíd, Sol, Fasola & Carranza, 2016; Oliver, Skipwith & Lee, 2014; Tallowin, Tamar, Meiri, Allison, Kraus, Richards & Oliver, 2018). In this study, we use two insular radiations of gecko lizards (Squamata: Gekkota), both potential ARs, to test the ecological axes of trait diversification, assess the strength of selection using tests of phenotypic convergence as a proxy, and investigate the relative importance of interspecific competition and environmental factors in shaping communities. With over 2,400 species, gekkotan lizards constitute a quarter of all squamates (lizards and snakes), with the bulk of species diversity being circumtropical. Further, geckos appear to be particularly well-represented on island systems, with numerous examples of long-distance transoceanic dispersal (Ali & Hedges, 2023; Greenbaum, Bauer, Jackman, Vences & Glaw, 2007; Raxworthy, Pearson, Zimkus, Reddy, Deo, Nussbaum & Ingram, 2008; Schwarz, Itescu, Antonopoulos, Gavriilidi, Tamar, Pafilis & Meiri, 2020; Skipwith, Bauer, Jackman & Sadlier, 2016; Skipwith, Bi & Oliver, 2019; Yoder & Nowak, 2006). In fact, Meiri (2019) notes that geckos are more likely to be insular than other squamates, with over a third (∼35%) being insular endemics. Diplodactyloids (families: Diplodactylidae, Carphodactylidae, and Pygopodidae) represent the predominate gecko lineage endemic to the southeastern Sahul shelf, largely replacing their sister taxon, the gekkonoids (families: Gekkonidae, Phyllodactylidae, Sphaerodactylidae, and Eublepharidae), which have a more cosmopolitan distribution west of Lydekker’s Line. Two lineages within the family Diplodactylidae have independently colonized the islands of New Caledonia and New Zealand during the Cenozoic (Nielsen, Bauer, Jackman, Hitchmough & Daugherty, 2011; Skipwith et al., 2016; Skipwith et al., 2019). In addition to being biodiversity hotspots, New Caledonia and New Zealand are remnants of the submerged continental fragment Zealandia (or Tasmantis), which separated from eastern Australia during the Campanian (∼80 mya). This region has perplexed biogeographers for decades as much of their respective biotas are most closely allied to Australian lineages, leading many authors to propose ancient vicariance due to continental drift as the primary force driving insular endemism (Bauer, 1990a; Espeland & Murienne, 2011; Giribet & Baker, 2019; Heads, 2008; Heads, 2010a; Heads, 2010b; Ladiges, 2008). However, the advent of molecular phylogenetics and dating methods has complicated this hypothesis as the vast majority of New Caledonian and New Zealand taxa investigated to date have yielded crown ages substantially younger than the widely accepted late Cretaceous eastern Gondwana-Zealandia rift (Chapple, Ritchie & Daugherty, 2009; Espeland, Johanson & Hovmoller, 2008; Espeland & Murienne, 2011; Giribet & Baker, 2019; Grandcolas, Murienne, Robillard, Desutter-Grandcolas, Jourdan, Guilbert & Deharveng, 2008; Michalak, Zhang & Renner, 2010; Murienne, Pellens, Budinoff, Wheeler & Grandcolas, 2008; Nielsen et al., 2011; Pratt, Morgan-Richards & Trewick, 2008; Skipwith et al., 2016; Skipwith et al., 2019; Smith, Sadlier, Bauer, Austin & Jackman, 2007; Trewick & Morgan-Richards, 2005; Trewick, Paterson & Campbell, 2007). Arguments for recent overwater dispersal are further supported by geological evidence that most of Zealandia may have been submerged, at least intermittently, until about 30 mya (Cluzel, Aitchison & Picard, 2001; Cluzel, Maurizot, Collot & Sevin, 2012). This may be particularly true for the smaller New Caledonia while there is some evidence that at least part of New Zealand remained aerial permitting lineage survival (Wallis & Jorge, 2018).

The two Zealandian diplodactylid assemblages are interesting in that they are both seen as candidate ARs, being ecomorphologically diverse with relatively young crown ages. Both lineages have independently evolved a number of traits and natural histories including gigantism, diurnality, adhesive distal scansors on the tail, and viviparity. Indeed, the largest extant and extinct geckos (> 250 mm SVL) are nested within the New Caledonian assemblage (Heinicke, Nielsen, Bauer, Kelly, Geneva, Daza, Keating & Gamble, 2023; Skipwith et al., 2016; Skipwith & Oliver, 2023). In general, biogeographers have assumed close phylogenetic ties between lineages present on both archipelagos and this is the case for the co-distributed lygosomine scincids lizards (Chapple et al., 2009; Smith et al., 2007). However, molecular evidence for diplodactylids illustrates that the Zealandian lineages are rather distantly related to one another, having diverged from different Australian lineages during the late Cretaceous or earliest Paleogene (Nielsen et al., 2011; Skipwith et al., 2016; Skipwith et al., 2019).

Convergent evolution has been well documented in several lineages, highlighting the deterministic aspect of phenotypic evolution in the face of selection (Bravo, Remsen & Brumfield, 2014; Burbrink, Chen, Myers, Brandley & Pyron, 2012; Zelditch, Ye, Mitchell & Swiderski, 2017). This result is illustrated in insular Greater Antillean *Anolis* lizards (terrestrial islands) and cichlid fishes of the Afrotropical great lakes (aquatic ‘islands’) (Arbour & Lopez-Fernandez, 2014; Mahler, Ingram, Revell & Losos, 2013; Mahler, Revell, Glor & Losos, 2010; McGee, Borstein, Meier, Marques, Mwaiko, Taabu, Kishe, O’Meara, Bruggmann, Excoffier & Seehausen, 2020; Patton, Harmon, del Rosario Castañeda, Frank, Donihue, Herrel & Losos, 2021; Ronco, Matschiner, Böhne, Boila, Büscher, El Taher, Indermaur, Malinsky, Ricci, Kahmen, Jentoft & Salzburger, 2021). In both systems, the repeated independent invasions of ‘islands’ has resulted in assemblages with replicated phenotypes and guilds. Previous studies, particularly those relying solely on morphology, suggested that the New Caledonian lineage is nested within the New Zealand clade or vice versa (Bauer, 1990a; Kluge, 1967a; Kluge, 1967b), contrasting with modern molecular inferences. This is not surprising given that both lineages are composed entirely of climbing forms preferring vegetation or rocky escarpments and are devoid of the terrestrial, saxicolous, and gramnicolous forms seen in their Australian relatives. Thus, it is reasonable to assume a broad overlap in morphospace between the two lineages consistent with selection-driven convergent evolution. Seemingly, like crown-giant *Anolis*, the New Caledonian lineage displays a pattern where larger species tend to live higher in the forest canopy than their smaller counterparts (Bauer & Sadlier, 2000). This pattern has been suggested for the distantly related gekkonid genus *Cyrtodactylus* (Oliver et al., 2014). Surprisingly, previous comparative studies have found mixed signals of either clade falling in line with the expectations of AR theory (Skipwith et al., 2016; Skipwith & Oliver, 2023). Namely, neither display exceptional speciation rates relative to other diplodactyloids or strong associations between speciation rate and trait diversification. Confoundingly, there is little evidence of early bursts in trait diversification and both lineages display highly idiosyncratic dynamics of integration amongst traits (Skipwith & Oliver, 2023).

Here, we investigate how habitat and ecological opportunity have impacted speciation dynamics and trait diversification in these two gecko radiations. Using a nearly complete phylogenomic estimate of the broader diplodactyloid radiation, we ask: (1) Is there convergent evolution between the New Caledonian and New Zealand lineages? (2) Are traits absent in mainland relatives such as distal caudal scansors, diurnality, and viviparity candidate key innovations driving speciation dynamics in Tasmantis forms? And (3) What is the relative importance of biotic and abiotic factors in shaping sympatry and character overlap in gecko communities? Contrasting these two superficially similar lineages presents an opportunity to quantify how selection and interspecific interactions influence character evolution.

## Methods

### Taxon Sampling

All sampling for this study is a subset of the datasets of Skipwith et al. (2019) and Skipwith and Oliver (2023), both focusing on the broader diplodactyloid radiation. From the phylogenomic estimate of Skipwith et al. (2019), we extracted the topologies of 36 New Caledonian and 18 New Zealand species, representing 95% and 90% of species, respectively. All described genera are present as well as all putative ecomorphs (see Morphological analyses). For tests of convergence, we incorporated the entire diplodactyloid tree from Skipwith et al. (2019).weused 298 voucher specimens for morphological analyses focusing on these two insular radiations and 993 specimens were used when in the context of the entire diplodactyloid radiation. All vouchers used in this study for morphological analysis were represented in the phylogeny (New Caledonia: 33 species; New Zealand: 18 species). In Skipwith and Oliver (2023) specific vouchers are clearly described and those were used for our morphometric analyses.

### Phylogenetic Analyses

This study relies entirely on the time-calibrated phylogenomic estimate of 180 diplodactyloids from Skipwith et al. (2019) using ∼4,200 ultraconserved elements (UCEs) representing ∼75% of recognized species diversity. Final in-target assemblies of UCEs are available on the Mendeley Data Platform (DOI:10.17632/zstmhjfkw5.1). We direct the reader to the supporting information of this manuscript and Skipwith et al. (2019) for details on phylogenetic reconstruction and NGS data assembly.

### Morphological Analyses, Disparity, and Testing Ecomorphology

Morphological analyses relied on a subset of the dataset used by Skipwith and Oliver (2023) and readers are directed there for details on data management and composition. In short, all species sampled here were represented by at least five adult specimens, with a special focus on males, where possible. A number of authors have noted that sexual dimorphism in trait proportions is rather modest in gekkotans when compared to iguanians, so we opted to keep species represented by females only in all downstream analyses (Kelly, 2015; Oliver, Ashman, Bank, Laver, Pratt, Tedeschi & Moritz, 2019; Skipwith & Oliver, 2023). All measurements were collected from the left side of the body using digital calipers by Phillip L. Skipwith.

All measurements were chosen due to their presumed ecomorphological relevance (Table S1). These were: body-size (SVL: snout-vent length), head length to retroarticular process (HLret), head width (HW), head depth (HD), trunk length (Trk), forearm length (Crus), tibia length (Tibia), inter- narial width (EN), nares-eye distance (NEYE), orbit diameter (EYE), eye-ear distance (EyeEar), interorbital distance (IE), fourth finger width (4FW), fourth finger length (4FL), fourth toe width (4TW), and fourth toe length (4TL). All comparative phylogenetic analyses used the species mean of each log-transformed trait. As these data are a subset of a previously published study, we relied on trait model selection outlined in Skipwith and Oliver (2023). All of the measurements used here are proxies for size, not shape, so we corrected for allometry by correcting all traits for body-size (SVL) using conventional (GLS) and phylogenetic least squares regression (PGLS) under Brownian motion (BM). We further corrected for element-specific allometry by correcting various cranial and limb measurements against the longest element of that region (i.e.-correcting HD for HLret, correcting 4TW for 4TL, etc.). This resulted in three multivariate datasets: (a) all traits regressed against SVL, (b) all cranial traits regressed against HLret, and (c) all digit and limb traits regressed against digit and limb length, respectively. We focused on the head and feet specifically because these are major axes of variation in geckos on the whole and clearly correspond to aspects of life history such as diet and locomotion (Garcia-Porta et al., 2016; Skipwith & Oliver, 2023; Vidal-García & Scott Keogh, 2017). The R package *phytools* was used for all PGLS analysis (Revell, 2012). To account for covariation amongst traits, the residuals of the three abovementioned datasets were reduced using both phylogenetic principal component analysis (pPCA) and conventional principal component analysis (cPCA).weopted to perform both forms of ordination in light of the concerns raised by Uyeda, Caetano and Pennell (2015) regarding model misspecification and bias in pPCA. However, Uyeda et al. (2015) notes that failing to account for phylogenetic signal in comparative analyses is also a form of model misspecification. Hence, we included pPCA results in downstream analyses where relevant, but only report on cPCA results if the two ordination methods yielded similar findings. Most of the following comparative analyses using residuals or eigenvectors from conventional or phylogenetic methods were estimated on the New Caledonian and New Zealand lineages separately. For a few analyses, namely tests of convergent evolution, the two were analyzed together with the entirety of the diplodactyloid radiation.

One of the important patterns observed amongst multiple AR examples is the fact that morphological and ecological diversity should evolve early during diversification before slowing down through time as niche-space contracts (Harmon, Losos, Jonathan Davies, Gillespie, Gittleman, Bryan Jennings, Kozak, McPeek, Moreno-Roark, Near, Purvis, Ricklefs, Schluter, Schulte Ii, Seehausen, Sidlauskas, Torres-Carvajal, Weir & Mooers, 2010). This pattern was tested in our clades by calculating disparity through time (DTT) from the mean-squared Euclidean distance between species, resulting in the morphological disparity index (MDI). Signatures of early burst are reflected as negative estimates of MDI (Harmon et al., 2010; Hipsley, Miles & Müller, 2014). We estimated DTT and MDI in the R package *Geiger* (Harmon, Weir, Brock, Glor & Challenger, 2008) on SVL alone and pPCA scores on cranial and limb traits independently. MDI was calculated from PCA scores estimated independently from the New Caledonian and New Zealand datasets which contrasts with MDI estimation in Skipwith and Oliver (2023) where MDI was estimated for datasets derived from the entire diplodactyloid lineage. This approach permits for the evaluation of lineage-specific primary axes of trait variation.

I used broad ecological guilds based largely on preferred substrate preference as a proxy for ecology. These are: (a) arboreal, living almost entirely on trees, (b) scansorial, generalized climbing on low vegetation, (c) and scansorial/rupicolous, living interchangeably of rocky surfaces and low vegetation. These categories were gleaned from the primary literature, detailed species accounts in specialized field guides, and databases (Bauer & Sadlier, 2000; Cogger, 2014; Meiri, 2024; Wilson & Swan, 2013).

### Adaptive Shifts and Convergent Evolution

We explicitly tested for convergent evolution and adaptive shifts along the diplodactyloid tree by modeling stabilizing selection. First, we classified putative ecomorphs into the three broad guilds mentioned above for the Zealandian lineages in addition to six guilds present in the broader Australasian diplodactyloid radiation. These additional guilds are: rupicolous (living primarily on rock faces), terrestrial (living primarily on the ground), scansorial/terrestrial (living on low vegetation and the ground), gramnicolous (living on grasses), limbless-terrestrial (snake-like pygopodids), and fossorial (living underground, *Aprasia*). These six guilds are not present in either of the insular lineages, but arboreal, scansorial and scansorial/rupicolous are present in the mainland lineages. Ancestral guild was reconstructed for all three diplodactyloid families using the 180 tip species tree of Skipwith et al. (2019). Ancestral states were reconstructed using Bayesian-based stochastic mapping which implements a Markov process to map shifts between discrete species states (Huelsenbeck, Nielsen & Bollback, 2003). Uncertainty in stochastic mapping reconstruction was accounted for by running 10,000 stochastic maps in order to calculate the posterior probability (PB) of each state. We tested three different models of discrete character evolution (equal-rates, symmetrical rates, all rates vary) using weighted Akiake information criterion (AICw) and ran all stochastic mapping analyses in the R package *phytools* (Revell, 2012; Revell, 2013).

A number of likelihood and Bayesian methods for automatically identifying adaptive shifts have been proposed and some have extended models of stabilizing selection to find convergent adaptive regimes amongst clades (Ingram & Mahler, 2013; Khabbazian, Kriebel, Rohe & Ane, 2016; Uyeda & Harmon, 2014). However, Adams and Collyer (2017) and Adams and Collyer (2019) note that some of these methods that do not require *a priori* regime identification, particularly *SURFACE* (Ingram & Mahler, 2013), are extremely susceptible to model misspecification and high typeweerror rates. Therefore, we tested adaptive shifts in morphospace without explicit tests of convergence across the limbed diplodactyloids (excluding the limbless pygopodids) using the likelihood-based *PhylogeneticEM* (Bastide, Ané, Robin & Mariadassou, 2018; Bastide, Mariadassou & Robin, 2016). PhylogeneticEM uses a simplified multivariate Ornstein-Uhlenbeck (OU) process, called the scalar OU (scOU), to automatically identify adaptive shifts. The scalar OU process offers advantages over multivariate OU models implemented in SURFACE (Ingram & Mahler, 2013) and *l1OU* (Khabbazian et al., 2016) in that it permits incidental correlations between traits.weestimated adaptive shifts separately for the three pPCA datasets on the limb-only diplodactyloid tree: all traits corrected for SVL, digit and limb traits corrected for digit and limb lengths, and cranial traits corrected for HLret. The total number and locations of estimated adaptive shifts were identified by the *params_process* function in PhylogeneticEM and model selection was done using linear estimator selection and lasso penalization.

Given the statistical shortcomings of SURFACE and other methods for identifying adaptive regimes without *a priori* groups,weopted to use methods that rely on designated discrete classifications. First, convergent evolution between the New Caledonian and New Zealand ecomorphs was explicitly tested using a derivation of phylogenetic ridge regression in the R package *RRphylo* (Castiglione, Tesone, Piccolo, Melchionna, Mondanaro, Serio, Di Febbraro & Raia, 2018).wetested the degree of phenotypic similarity between *a priori* diplodactyloid ecomorphs using the *search.conv* function in RRphylo (Castiglione, Serio, Tamagnini, Melchionna, Mondanaro, Di Febbraro, Profico, Piras, Barattolo & Raia, 2019). This method treats the multivariate data for each species as a vector, where θ represents the angle between each species pair. The degree of phenotypic similarity between this pair is calculated as the cosine of θ, with a value range spanning 0-180 degrees. A low value of θ indicates stronger phenotypic similarity as 0° is approached and increasing dissimilarity when approaching 180°. In short, similarity suggesting convergence between entire clades or species grouped under *a priori* assignments is determined when θ is smaller than expected given phylogenetic distance under Brownian motion (BM).wefirst implemented a state-agnostic run of search.conv where the algorithm automatically searches for convergence across the phylogeny without *a priori* ecomorph classifications. We then ran the state-informed version where each ecomorph (arboreal, scansorial, and scansorial/rupicolous) was tested separately for (1) all limbed diplodactyloids and (2) just the New Caledonian and New Zealand lineages. This design allowed me to test if convergence was present between all lineages for a given ecomorph or if convergence was exclusive to the insular lineages.

The strength of convergence for ecomorphs identified by RRphylo was tested with the Wheatsheaf index (*w*) in the R package *Windex* (Arbuckle, Bennett & Speed, 2014; Arbuckle & Minter, 2015). Here, the potentially convergent lineages for a single ecomorph are treated as focal taxa *a priori* and *w* is calculated from the similarity of focal taxa and contrasting this to their dissimilarity to non-focal taxa. This is accomplished through the construction of a phenotypic distance matrix where the Euclidean distance (*d_ij_*) between species*e*and *j* is corrected for phylogenetic distance by dividing by 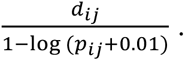 Here, *p*_ij_ is the shared proportional distance between species *i* and *j*. Using *d_ij_*, *w* is the quotient of mean pairwise comparisons of all species (*d_a_*) divided by the mean pairwise comparisons of all focal species (*d_f_*). As *w* increases, the strength of convergence increases, suggesting greater overlap in morphospace than expected. The Wheatsheaf index was estimated using the same analytical design and datasets as RRphylo for each proposed ecomorph in Windex with 1,000 bootstrap replicates. From the bootstrap sampling, the 95% confidence level for *w* was obtained through jackknifing along with the reported p-value.

### ABIOTIC AND BIOTIC FACTORS DRIVING SPECIATION

Theory suggests that ecologically similar species should not overlap spatially in order to avoid interspecific competition. However, many candidate ARs are systems where closely related taxa coexist with minimal competition, presumably due to distinct phenotypes and nonoverlapping niches (Pfennig & Pfennig, 2020; Schluter, 2000a). Distinguishing how character displacement due to divergent selection and environmental factors impact community composition has been made possible through recent advances in computational methods. We test for the presence of character displacement independently in the New Caledonian and New Zealand diplodactylids using a recently developed Bayesian parametric model that incorporates both trait data and biogeographic spatial data in the Julia package *TAPESTREE* (Quintero & Landis, 2019). The Bayesian *TRIBE* model in TAPESTREE implements an event-based model that quantifies how lineages evolve independently of one another and how trait dynamics and geographic range impact one another. Further, the TRIBE model quantifies the relative importance of pre- and postsympatric niche divergence (Quintero & Landis, 2019). This probabilistic model differs from some other explicit models of character displacement in that it simultaneously models how trait values for all species in a given area impact colonization and extirpation in that area as well how trait values respond to those of sympatric species. There are three parameters estimated here that are of particular importance in sympatry: sympatric trait evolution (ω_x_), the effect of biotic interactions on extirpation (ω_0_), and the effect biotic interactions on colonization (ω_1_). When any of these parameters has in their credible interval the value of 0, there is no inferred biogeographical effect on the trait for species sharing space. Lastly, environmental filtering is implied when ω_0_ < 0 or ω_1_ > 0 whereas competitive exclusion is implied when ω_0_ > 0 or ω_1_ < 0. We ran the TRIBE model for SVL and each pPCA and cPCA score for cranial and limb traits separately for the New Caledonian and New Zealand taxa for 600,000 generations with a burnin of 50,000 generations. Importantly, these PCA scores were calculated from ordinations of each Zealandian clade separately, not as part of the larger diplodactyloid radiation. Therefore, they are distinct from those used in all of the abovementioned analyses on adaptative diversification and convergence, highlighting differing signals of lineage-specific axes of trait variation. Each TRIBE analysis was run twice and convergence was assessed using the R package Coda (Plummer, Best, Cowles & Vines, 2006) that checked for Gelman diagnostic tests and estimated sample sizes (ESS ≥ 200 taken as good mixing of MCMC chain), and also visually from *Tracer* (Drummond, Suchard, Xie & Rambaut, 2012)

The TRIBE model requires both trait data and range data and we used a combination of proposed ecoregion and floral province classifications to discretize the ranges of Zealandian geckos. The ecoregion boundaries of New Zealand are fairly well characterized in a global context andweused the terrestrial ecoregion classifications proposed by Dinerstein, Olson, Joshi, Vynne, Burgess, Wikramanayake, Hahn, Palminteri, Hedao, Noss, Hansen, Locke, Ellis, Jones, Barber, Hayes, Kormos, Martin, Crist, Sechrest, Price, Baillie, Weeden, Suckling, Davis, Sizer, Moore, Thau, Birch, Potapov, Turubanova, Tyukavina, de Souza, Pintea, Brito, Llewellyn, Miller, Patzelt, Ghazanfar, Timberlake, Klöser, Shennan-Farpón, Kindt, Lillesø, van Breugel, Graudal, Voge, Al-Shammari and Saleem (2017) with four states. These were: the South Island temperate forests, South Island grasslands, North Island temperate forests, and Rakiura Island temperate forests. Unfortunately, ecoregion classifications for New Caledonia are somewhat coarse in comparison.

Thus,wechose to use three broad groupings based on the primary vegetation type and supported by a number of biogeographers (Bauer & Sadlier, 2000; Jaffré, Bouchet & Veillon, 1998; Murienne, 2009). These were: maquis, sclerophyll forests, and humid forests.

To summarize the relative contribution of abiotic and biotic factors on speciation in Zealandian diplodactylids,weused phylogenetic path analysis (PPA) (Gonzalez-Voyer & von Hardenberg, 2014; Hardenberg & Gonzalez-Voyer, 2013). PPA is class of multiple phylogenetic regressions that incorporates phylogenetic relationships into models of trait evolution, allowing investigators to test for direct and indirect causal relationships between traits.weused the R package *Phylopath* (van der Bijl, 2018) to evaluate a series of nested models on the relationships between speciation rate, phenotypic evolutionary rate, climate niche evolution, and habitat. Rates of speciation and trait evolution were estimated from tip rates which are particularly reliable estimators for the former (Title & Rabosky, 2019). Studies investigating speciation rates are particularly sensitive to tree size, particularly regarding parameter estimation and model misspecification underlying small phylogenies. In order to alleviate some of these issues, we used the log-transformed speciation tip rates estimated with BAMM (Rabosky, 2014; Rabosky, Grundler, Anderson, Title, Shi, Brown, Huang & Larson, 2014; Rabosky et al., 2013) from Skipwith and Oliver (2023) on the larger, taxonomically complete tree 241 species tree of all diplodactyloids simulated with the stochastic polytomy resolver TACT (Chang, Rabosky & Alfaro, 2019). The speciation tip rates were extracted for just the New Caledonian and New Zealand lineages. To complement tip rates estimated with BAMM, we used branch rates estimated with ClaDS from Skipwith and Oliver (2023) with constant extinction (ClaDS1) (Maliet, Hartig & Morlon, 2019). Trait data included eight variables, six continuous and two discrete. These were: speciation rates (continuous: SpecRates), rates of body-size evolution (continuous: SVLrates), rates of head shape evolution (continuous: HeadRates), rates of limb evolution (continuous: LimbRates), rates of climatic niche evolution (continuous: ClimRates), rates of elevation occupancy evolution (continuous: ElevRates), ecology (discrete: Eco), and habitat (discrete: Hab). Cranial morphology and limb morphology were based on the same PC scores for each clade as described for analyses of disparity and convergent evolution. Minimum and maximum elevation data was gleaned from species descriptions or from SquamBase, a life history database for virtually all squamate species compiled by Meiri (2024). Range data for each species of New Caledonian and New Zealand diplodactylid was extracted from observation records on the Global Biodiversity Information Facility (GBIF, available at http://www.gbif.org) using the R package *rgbif* v3.8.2 (Chamberlain, 2017). Supplemental data was also cleaned from the Atlas of Living Australia (ALA, https://www.ala.org.au/, (Belbin, Wallis, Hobern & Zerger, 2021)) using the R package *galah* v2.1.2 (Westgate, 2025). We then generated shapefiles for each species observation in order to obtain mean estimates of the 19 climatic variables from WorldClim v2.1 (available at http://www.worldclim.org) at a resolution of 2.5 km. As with the morphological data, covariation amongst climate and elevation data were accounted for with their own PCAs. We then used the RRphylo function in *RRphylo* to estimate tip rates of trait evolution on SVL and on the pPCA scores (top ∼90% of variance explained) for head traits, limb traits, climate, and elevation. PPA with *PHYLOPATH* only allows for binary discrete characters, sowebinarized habitat (Hab: forest vs. maquis/savannah) and ecology (Eco: arboreal vs. scansorial). A total of twenty-eight increasingly complex models were constructed in *PHYLOPATH* to assess the dependence of speciation rates (SpecRates) on morphology (SVLrates, HeadRates, and LimbRates), life-history (Eco and Hab), and abiotic factors (ClimRates and ElevRates). These models included single direct effects (i.e.-SpecRates influenced by SVLrates), multiple direct effects (i.e.- SpecRates influenced by SVLrates and ClimRates), and multiple indirect effects (i.e.- SpecRates influenced by SVLrates which are dependent on ClimRates). In addition to contrasting the Zealandian assemblages,wetested whether similar processes govern diversification across all limbed diplodactyloids. In these analysesweused the same continuous characters (SpecRates, SVLrates, HeadRates, LimbRates, ClimRates, and ElevRates) for 129 species of limbed diplodactyloid and five binarized discrete characters. These were: biogeography (BioGeo: mainland vs. insular), reproductive mode (Repro: oviparous vs. viviparous), diel pattern (Activity: nocturnal vs. diurnal/cathemeral), adhesive digital scansors (Pads: padded vs. padless), and terminal caudal scansors (TailPads: padded vs. padless). Further, many of these characters are distinct and appear independently in the Zealandian lineages (viviparity, diurnality, and adhesive tail tips), presenting themselves as potential key innovations influencing speciation. We tested forty-four alternative models of indirect and direct biotic and biotic factors (Figs. S1 and S2). *Phylopath* analysis used Pagel’s lambda (λ) constrained between 0 and 1 for regressions of continuous variables, penalized likelihood of the logistic regression of binary variables (MPLE), and Fisher’s C statistic to evaluate model fit (Gonzalez-Voyer & von Hardenberg, 2014). Model fit was further assessed using the corrected *C*-statistic criterion (CICc), where a ΔCIC < 2 indicated similar support for different models. In cases where ΔCIC > 2, the best model was verified with 500 bootstrap replicates whereas when ΔCIC < 2, models were averaged.

## Results

### Unexceptional Disparity in Insular Lineages

Details on the variance explained by individual cranial and limb traits in pPCAs and cPCAs are outlined in the supplement (Table S2). Analysis of morphological disparity across body-size (SVL), cranial dimensions, and limb proportions for the each of the Zealandian lineages yielded largely similar patterns. Neither lineage exhibited exceptional disparity relative to their respective ages when examining cranial or limb dimensions (Table 1 and Table S3). However, the New Caledonian clade did display high disparity acquired early in the radiation when focusing on body-size alone (MDI = -0.26, p = < 0.01).

**Table 1:**
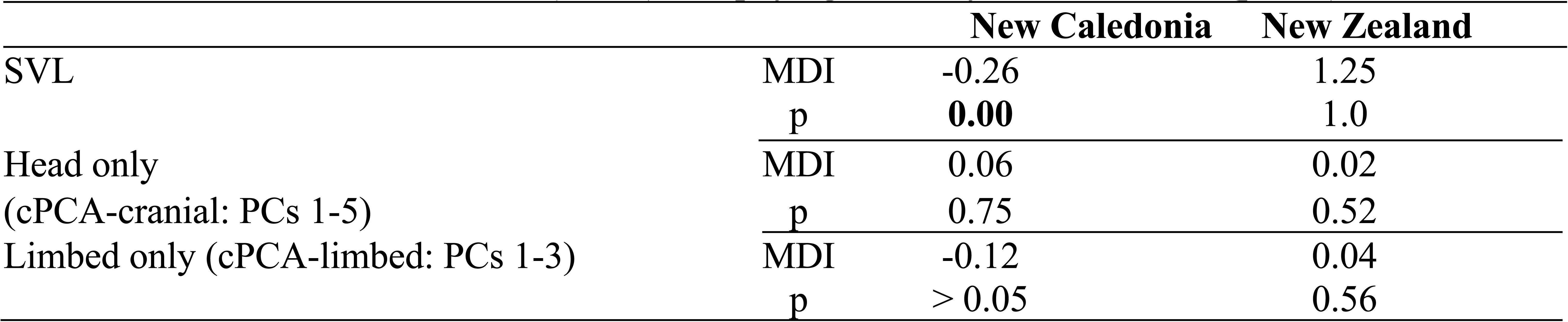
Mean disparity indices (MDI) for both of the Zealandian clades with p-values from 1,000 simulations under BM. Bold p-values indicate significance. Each clade was analyzed seperately. All estimates are for conventional PCA (cPCA), not phylogenetically corrected PCA (pPCA).

### Adaptive Shifts and Convergent Evolution

Model selection for models of discrete character evolution with AICw selected the equal rates (ER) model for the nine designated states. Ancestral state reconstruction using stochastic mapping on guild recovered a rupicolous origin for crown diplodactyloids (PB = 0.58), diplodactylids (PB = 0.53), the NC-*Pseudothecadactylus* clade (PB = 0.54), NZ-core Australia (PB = 0.43), and core Australia (PB = 0.42) with high confidence (Fig. 1). These reconstructions suggest that the respective ancestors of the NC and NZ clades differed in ecology with the former receiving unambiguous support for a scansorial ecology (PB = 1.0) while the NZ clade was received mixed support for a arboreal or scansorial origin (PB: arboreal = 0.32, scansorial = 0.35). Arboreal and scansorial/rupicolous guilds appear to have evolved independently multiple times in both the NC and NZ clades. Further, these same guilds have also evolved independently in the core Australian lineage with many transitions between arboreal, scansorial, and scansorial/rupicolous guilds in the plesiomorphic *Amalosia*-*Oedura* grade. See supplement for details on transition frequencies.

**Figure 1:**
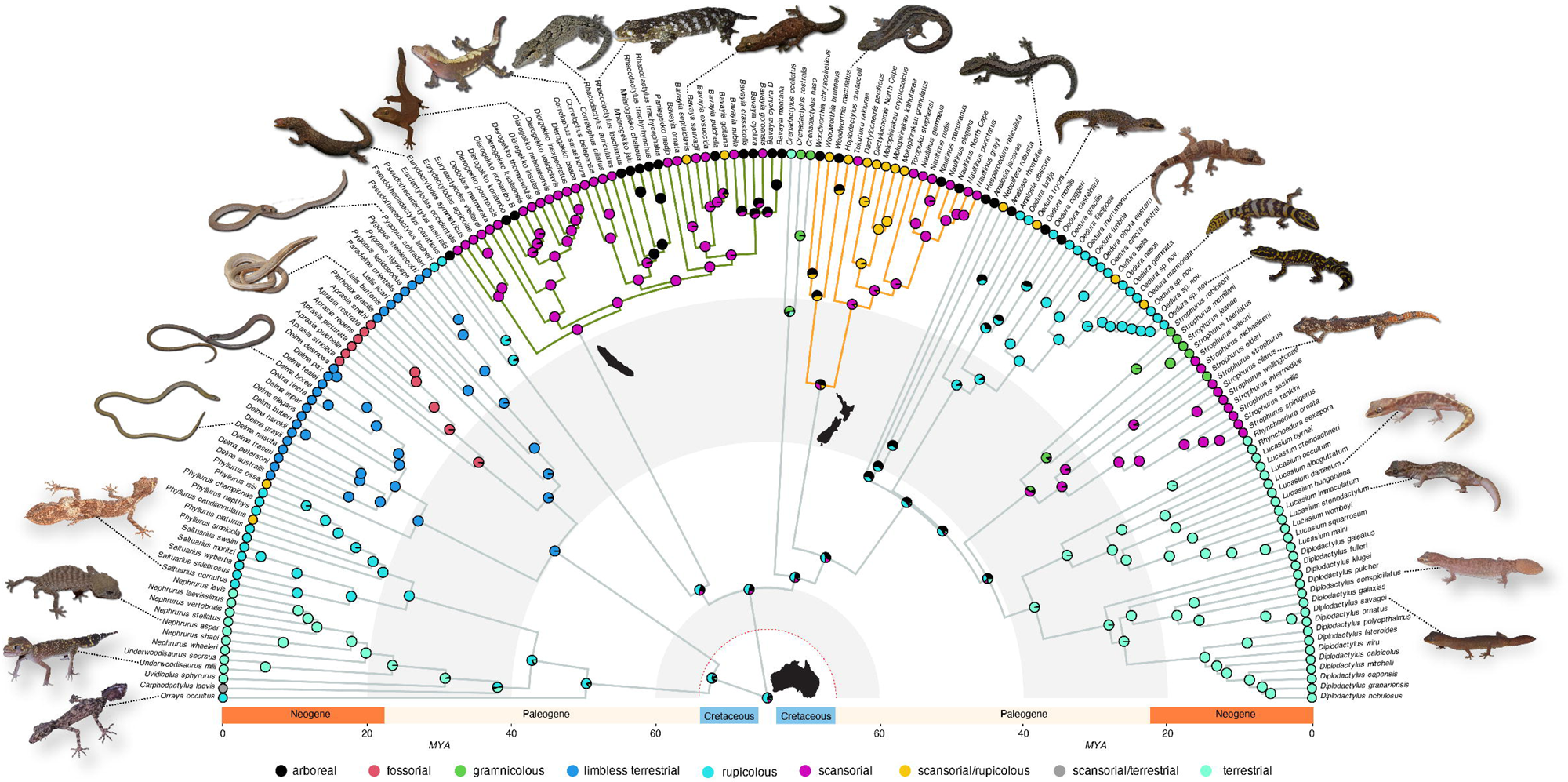
Timetree of 180 diplodactyloid species from Skipwith et al. (2019) with stochastically-mapped reconstructions of ecological guild. The New Caledonian clade is highlighted with green branches while the New Zealand clade is orange. Red dotted line at 66 mya denotes the KpG boundary. Silhouettes of Australia, New Caledonia, and New Zealand are not to scale All gecko images were taken by Phillip L. Skipwith and Paul M. Oliver.

Analysis of adaptive shifts with PhylogeneticEM identified shifts in all three pPCA datasets, though they differed somewhat in number, location, and magnitude between all traits cranial traits (k = 7), and limb traits (k = 6). While the cranial and limb datasets failed to recover shifts at identical locations on the phylogeny, they largely agreed by proximity (Fig. 2 A and B). For instance, both agreed on there being shits around the *Correlophus*-*Mniarogekko*-*Rhacodactylus* clade and at the base of or nested within the NZ clade. All shifts recovered by PhylogeneticEM correspond to clades with strong evidence of guild transitions in ancestral state reconstructions. See Figs. S3-S8 for detailed Phylogenetic EM plots for both cPCA and pPCA datasets.

**Figure 2:**
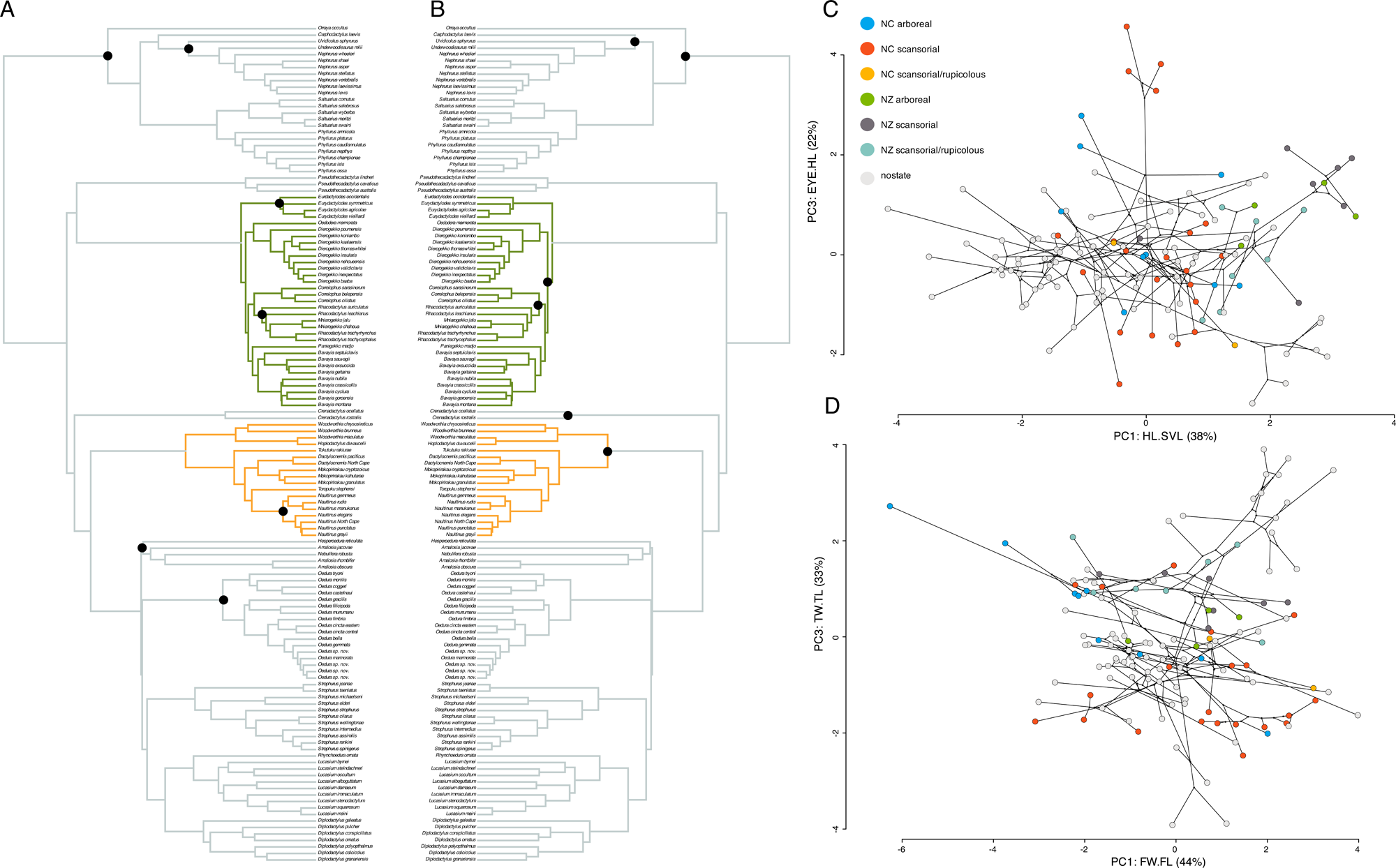
Timetree of 129 limbed diplodactyloids (pygopodids excluded) used in PhylogeneticEM analyses with ordination using cPCA. A) Cranial traits only. Black node circles indicate clades which experienced adaptive shifts. B) Limb traits only. Black node circles indicate clades which experienced adaptive shifts. C) Phylomorphospace plot of conventional PCA of cranial traits, colored by ecological guild for the NC and NZ clades only. D) Phylomorphospace plot of conventional PCA of limb traits, colored by ecological guild for the NC and NZ clades only.

Signatures of convergent evolution were identified for all three guilds replicated across the NC and NZ assemblages. Tests of convergence using phylogenetic ridge regression in RRphylo differed depending on whether *a priori* state was taken into account. State agnostic analyses failed to find convergence between the NC and NZ clades regardless of meristic data (all, cranial, or limb). However, state-informed analyses recovered strong support for convergence between all NC and NZ guilds for all meristic datasets except for the cranial scansorial/rupicolous dataset. The two metrics returned by search.conv, ang.state and ang.state.time, represent to the average θ_real_ between a species pair and the θ_real_ /divergence time, respectively. Castiglione et al. (2019) note that ang.state.time is the more relevant of the two metrics to note as the phylogeny sampled here is largely complete. Virtually all comparisons of NC and NZ guilds yielded significant estimates of ang.state.time (p < 0.01), suggesting that convergence and not parallel evolution is more prominent in this this system (Table 2). Additionally, search.conv recovers signatures of convergence between a number of insular guilds and their counterparts in the core Australian diplodactylid assemblage, particularly within the NC clade when considering head and limb traits (supp. Table 2). However, analysis of the strength of convergent evolution with *w* suggests that evidence of convergence is strong when examining cranial and limb proportions. Here, Windex found strong support for convergence (p ≤ 0.05) for all three guilds across all three datasets except for the arboreal and scansorial comparisons of all traits corrected for SVL (Table 3). These same analyses yielded largely identical patterns when using pPCA scores (see Tables S4 and S5) and will not be discussed further.

**Table 2:**
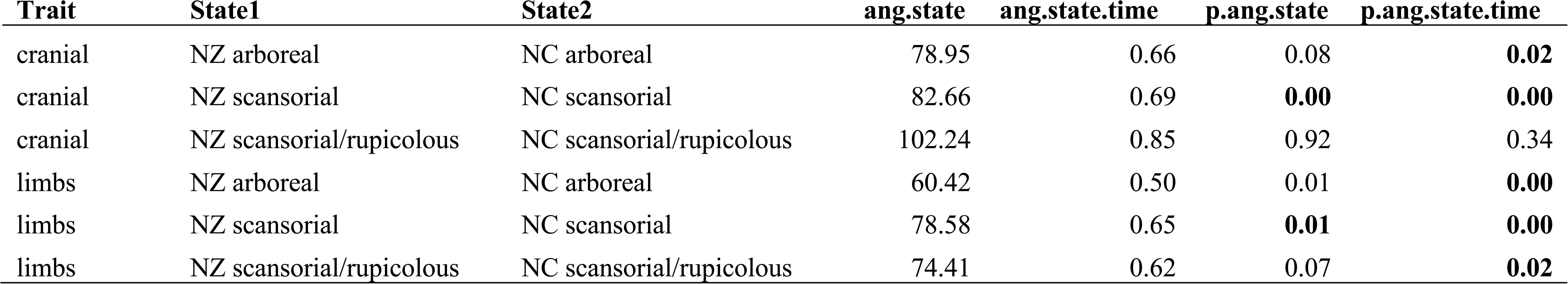
Parameter estimates of RRphylo tests of convergence (search.conv). Contrasted are the cranial and limb proportion datasets for each the three *a-priori* ecological guilds. Of particular value are the parameters *<u>ang.state.time</u>* and *p.ang.state.time*, which are stronger indicators of the presence of convergent evolution. All estimates are for conventional PCA (cPCA), not phylogenetically corrected PCA (pPCA).

**Table 3:** Parameter estimates of the Wheatsheaf index (*w*) from Windex. *w* is a metric for assessing the strength of convergence, no the presence of convergence (see results from RRphylo). Contrasted are the cranial and limb proportion datasets for each the three *a-priori* ecological guilds. All estimates are for conventional PCA (cPCA), not phylogenetically corrected PCA (pPCA).

| Analysis | State | $w$ | low95 | up95 | P |
| --- | --- | --- | --- | --- | --- |
| all traits corrected for SVL | arboreal | 0.72 | 0.69 | 0.73 | 0.63 |
| all traits corrected for SVL | scansorial | 0.67 | 0.66 | 0.69 | 0.26 |
| all traits corrected for SVL | scansorial/rupicolous | 1.04 | 1.03 | 1.07 | **0.01** |
| cranial | arboreal | 1.08 | 1.06 | 1.12 | **0.00** |
| cranial | scansorial | 0.76 | 0.75 | 0.77 | **0.01** |
| cranial | scansorial/rupicolous | 1.29 | 1.27 | 1.34 | **0.00** |
| limbs | arboreal | 0.76 | 0.74 | 0.78 | 0.44 |
| limbs | scansorial | 0.64 | 0.63 | 0.65 | 0.56 |
| limbs | scansorial/rupicolous | 1.05 | 1.04 | 1.09 | **0.02** |

### Abiotic Factors Drives Speciation

The TRIBE module recovered no signature of character displacement (P(ω_x_ <0)< 0.05) across any of the datasets for either Zealandian clade (Fig. 3) However, for the first principal component for limbs in the New Zealand clade we found evidence of morphological convergence in sympatry (P(ω_x_ >0)= 0.95). Both clades were further found to have low colonization and extirpation rates (ω_0_ < 0 and (ω_1_ < 0), across principal components. However, the 95% HPDs of ω_1_ show that phenotypically similar taxa do not colonize currently occupied niches. Meanwhile, sympatry has no effect on extirpation rates (P(ω_0_ <0)< 0.05).

**Figure 3:**
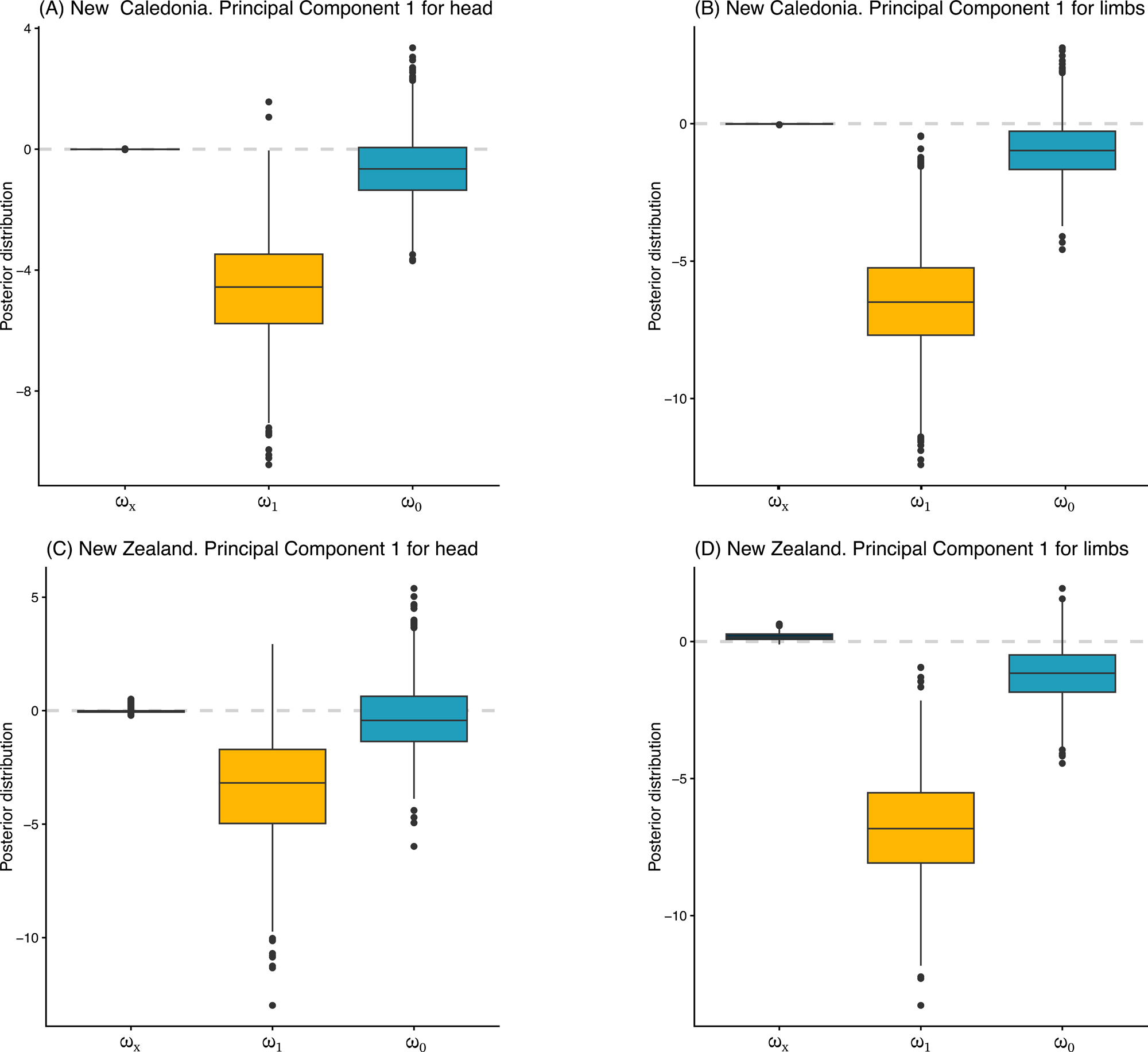
Analysis of character displacement. 95% HPDs of cPCA1 for A) New Caledonian cranial, B) New Caledonia limb, C) New Zealand cranial, and D) New Zealand limb. ω variables are: sympatric trait evolution (ω_x_), the effect of biotic interactions on extirpation (ω_0_), and the effect biotic interactions on colonization (ω_1_).

Comparing the twenty-four models contrasting direct and indirect effects using PPA where both clades seem to be best fit by an indirect model (model twenty-one, see supp.) where ClimRates -> LimbRates -> SpecRates (Fig. 4 B and C). Here, rates of climate niche evolution have strong positive effects on the rates of limb dimension evolution which in turn negatively impacts speciation rate. When expanded to the entire diplodactyloid radiation, PPA model selection yielded accordingly more complex scenarios. A model averaged across three alternatives recovered a scenario where biogeography negatively impacts ClimRates and ElevaRates, ElevaRates negatively impacts ClimRates, and both ClimRates and ElevaRates positively impacting speciation rate (Fig. 4 A). It should be noted that PPA using the pPCA generally differed in model selection for the NC, NZ, and all limbed diplodactyloids from those selected for the cPCA (see supporting information). However, both cPCA and pPCA PPA analyses selected models that strongly favored the effects of climate on trait diversification which in turn influenced speciation rates for both insular lineages.

**Figure 4:**
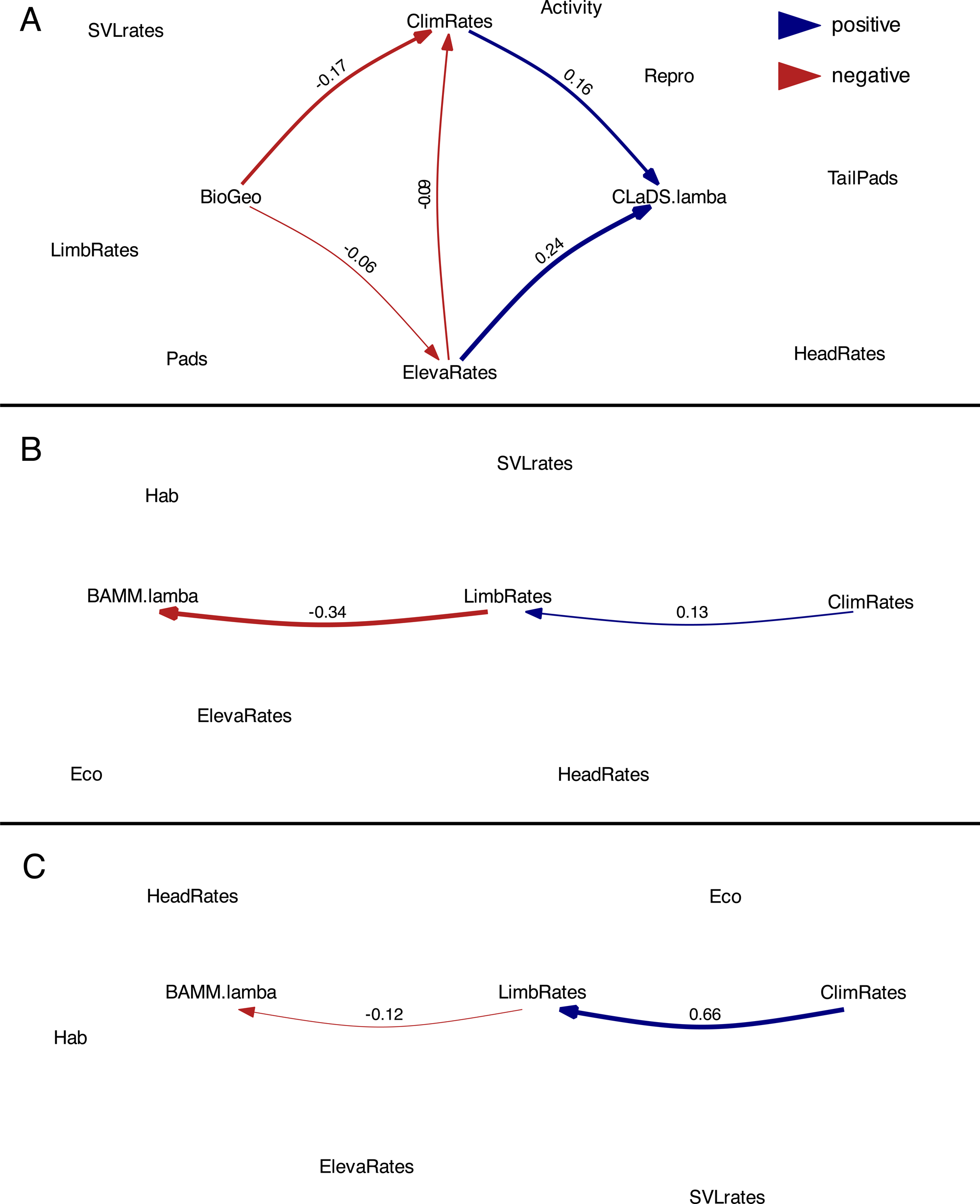
Output of PhyloPath Analyses on A) The 129 species tree for limbed diplodactyloids, B) the New Caledonian clade alone, and C) the New Zealand clade alone. Red and blue arrows indicate the magnitude and direction of variable predictive power.

## Discussion

### Ecomorphological Diversification and Convergence

In congruence with the findings of Skipwith and Oliver (2023), we found compelling evidence for unexceptional speciation rates and phenotypic disparity in Tasmantis diplodactyloids. This opposes the expectations of AR theory, which is surprising given that the crown ages of both lineages are relatively young when considering high species richness and the degree of ecomorphological variation present. This is particularly marked in the New Caledonian assemblage where giant arboreal forms such as *Rhacodactylus leachianus* (280 mm SVL) and *R. trachyrhynchus* (190 mm SVL) can occur in sympatry with small-bodied, scansorial lineages such as *Dierogekko* (38 – 45 mm SVL) and *Bavayia sauvagii* group (50-72 mm SVL) which primarily occur on low vegetation and rocky surfaces. In some communities, particularly on the southern half of Grande Terre (New Caledonia’s primary island), these forms may occur with medium-large scansorial forms (100 – 150 mm SVL) such as *Correlophus*, *Mniarogekko*, and *Rhacodactylus auriculatus*. *R. auriculatus* is notable in that it seems to bear dental adaptations for hypercarnivory broadly absent in geckos and may be a major predator of vertebrates (Bauer & Russell, 1990; Bauer & Sadlier, 1994; Bauer & Sadlier, 2000; Snyder, Snyder & Bauer, 2010). The NZ clade shows similar patterns where the giant *Hoplodactylus duvaucelii* (160 mm SVL) can cooccur with smaller *Woodworthia* (60 – 70 mm SVL) and *Naultinus,* (70 - 90 mm SVL). Of the traits examined, only body-size within the NC clade displays disparity in line with the expectations of AR, a finding also recovered by Skipwith and Oliver (2023) using a distinct ordination scheme. The overall failure to find signals of early burst in morphology in the Tasmantis forms suggests that either selection has not historically driven wholesale phenotypic diversification or that trait diversification is limited by lability. These explanations are made more viable as explicit tests of covariance and coevolution amongst traits suggest that traits are largely integrated, and selection may only liberate isolated traits to diverge in specific lineages (Skipwith & Oliver, 2023).

Ancestral state reconstruction, quantification of phenotypic shifts, and tests of convergent evolution strongly suggest that the NC and NZ clade initially began as ecologically and morphologically distinct with repeated evolution of similar phenotypes. In the case of the former, there appears to be some level of phenotypic conservatism as *Pseudothecadactylus* is superficially so similar to the NC giants (*Rhacodactylus*, *Correlophus*, and *Mniarogekko*) that Bauer (1990b) lumped the four genera into *Rhacodactylus* based on osteology. In contrast, the NZ clade is seemingly more complex as their closest relatives, *Crenadactylus* and the core Australian forms, are radically different in morphology and largely do not overlap in morphospace (see supp. Fig 1). Notably, there are multiple transitions to arboreality from scansorial lineages and ecomorphology suggests that these tree-dwelling forms have wider digits relative to length. Where convergence was strongly recovered for the arboreal and scansorial guilds across the cranial and limb datasets, it appears that arboreality is particularly complex. Intuitively, the arboreal giants of *Rhacodactylus* and *Mniarogekko* are unlikely to be under the same selective pressures as small-bodied arboreal forms such as *Dierogekko* (NC) and *Naultinus* (NZ). Rather, it appears that these large forms are convergent with another lineage of NC geckos, the intermediate-sized *Bavayia cyclura* group, whereas some arboreal *Dierogekko* and *Naultinus* converge in overall proportions (Fig. 2D). In the case of the Tasmantis scansorial and scansorial/rupicolous forms, it appears that digits are longer and narrower than in arboreal forms. Again, there are repeated transitions to these guilds in both lineages, but, with the exception of the giant *Hoplodactylus duvaucelii*, these are largely restricted to small-bodied forms inhabiting low vegetation.

While convergence was detected between the NC and NZ arboreal and scansorial forms, it was not indicated for the scansorial/rupicolous forms when examining the head. This may be attributable to a low degree of morphological specialization in the scansorial/rupicolous guild on their respective archipelagos. On New Caledonia, scansorial/rupicolous forms (as defined here) are restricted to two species in the *Bavayia sauvagii* complex whereas New Zealand’s representatives from the same guild are mostly clustered in the *Dactylocnemis*/*Mokopirirakau* clade. The *Bavayia sauvagii* clade, for the most part, bear relatively wide scansors while the *Dactylocnemis*/*Mokopirirakau* clade is nested within a lineage with comparatively narrow scansors (Nielsen et al., 2011). This pattern suggests that phylogenetic inertia may be partly responsible in both clades and that the biomechanical requirements for traversing both vegetation and rock surfaces are indistinguishable from those pertaining to predominately navigating the latter. Explicitly accounting for stabilizing selection with PhylogeneticEM reveals that numerous shifts are clustered either at the crowns of either Tasmantis lineage or nested within, though there is little consensus when contrasting conventional (cPCA) and phylogenetic (pPCA) ordination methods or body region (cranial vs. limbs). However, it seems that shifts in cranial morphology are less correlated with locomotion and more with diet and sensory perception. For example, both *Eurydactylodes* (NC) and *Naultinus* (NZ) are diurnal and are recovered in all analyses of the head to have experienced adaptive shifts in cranial proportions. Both genera have small eyes relative to head length, possibly an adaptation to diurnality (Hall, 2008; Werner, 1969), but do not overlap with one another in morphospace for other cranial traits or limb proportions. Notably, *Eurydactylodes* is aberrant in comparison to most geckos, being small-bodied with short limbs and digits. The proportions of *Eurydactylodes* are similar to the unrelated Greater Antillean *Anolis* twig ecomorphs, which are also relatively slow moving forms specialized for clinging to thin, low branches (Losos, 1990; Losos, 2009; Losos, Jackman, Larson, de Queiroz & Rodriguez-Schettino, 1998). However, the failure to recover shifts in adaptive optima for all lineages that transitioned guilds may be an artefact of large shifts to new optima having occurred early in diversification while subsequent shifts were of comparatively low magnitude.

### Abiotic and Biotic Factors Driving Diversification

When assessing character displacement with TRIBE, Quintero and Landis (2019) describe potential interpretation of their parameters as follows: when ω_x_ < 0 sympatry drives traits to diverge and when ω_x_ > 0, traits converge; when ω_0_ < 0 phenotypically similar taxa are less likely to be extirpated in sympatry while the inverse is true when ω_0_ > 0; and when ω_1_ < 0 phenotypically similar species have low rates of colonization and the inverse is true when ω_1_ > 0. Despite exhibiting extensive ecomorphological diversity in sympatry, neither Tasmantis clade displays evidence of character displacement (ω_x_ < 0). Rather, both lineages display some degree of character convergence in sympatry (ω_x_ > 0), and this is highly probable in the limbs of the New Zealand clade. Additionally, phenotypically similar taxa were estimated to have lower colonization rates (ω_1_) and lower extirpation rates (ω_0_) in sympatry. Taken together, this suggests that both competitive exclusion and niche filtering play a role in community assembly. The HPDs of ω_0_ overlap with 0 across datasets, suggesting that sympatry, in fact, may have no significant effect on extirpation. The effect is mostly shown in colonization rates (ω_1_), where taxa were prevented from colonizing currently occupied niches with phenotypically similar taxa. These TRIBE analyses paint an interesting picture on the formation of species assemblages on both archipelagos where biotic factors or, at the very least, interactions between closely related species have not played a major role historically.

The fact that phenotypes tend to converge in sympatry in diplodactyloids is surprising given the prevailing view that interpopulation competition may drive divergent selection in sympatry in ARs (Gillespie et al., 2020; Schluter, 2000a). Given that the TRIBE model is sensitive to phylogenetic uncertainty, the exclusion of extinct taxa, restricted to univariate traits, and assumes that biotic interactions are uniform across all lineages and time, it is tempting to dismiss this finding. However, these findings are more compelling when viewed in combination with PPA. While not analytically analogous to TRIBE, multiple phylogenetic regressions highlight that the tempo of climatic and elevational niche evolution have a much greater influence on speciation rates than morphometric traits or proposed key innovations (diel pattern, digit morphology, caudal morphology, and reproductive mode). Taken together, these findings suggest that sympatry is driven by populations that have preexisting adaptations and/or constraints that may have evolved in allopatry. The interplay between elevational and climatic niche evolution may be driven by adaptations that facilitate or hamper the ability of gecko populations to colonize new regions or adapt to changing environs. These data, however, cannot tease apart the relative importance of intrinsic physiological adaptions in geckos to climatic variables or adaptations to floral communities that are driven by these same variables. It is conceivable that gecko lineages adapted to specific climatic profiles or to the flora that are managed by the same environmental variables would be phenotypically similar and therefore survive in similar habitats. Here, secondary sympatry through rare migration events sets the stage for phenotypically convergent taxa to experience competitive exclusion, where subsequent extirpation is infrequent. This is consistent with niche filtering and competitive exclusion being identified with weak and moderate support, respectively, in the TRIBE analyses.

Our individual assessments of the NC and NZ lineages paint a picture where sympatry in these complex radiations is driven by adaptations to the environment (climate and elevation) and, possibly, biotic interactions with other aspects of their communities, not interactions between competing populations of gecko. While each of the Tasmantis lineages occupy distinct climatic niches from one another and all other limbed diplodactyloids, PPA finds that largely similar patterns are found across the broader diplodactyloid assemblage. Here, biogeography (insular vs. mainland) affects climatic and elevational niche diversification which, in turn, drive speciation. This suggests that while each clade of limbed diplodactyloid diversifies along distinct ecomorphological axes, there is relative conservatism in the importance of climate and topography in determining where gecko lineages occur and the spatial distribution of ecomorphs. It may also be indicative that speciation and ecological divergence is driven by allopatric processes with, potentially little opportunity for the erosion of nascent species boundaries through introgression. Though, clade-wide population genetic data would be needed to test that hypothesis. This may be a pattern that is broadly true across Gekkota, but the restricted geographic range of diplodactyloids in comparison to their sister taxon, the gekkonoids, suggests that this is a far more complicated picture.

Identifying key innovations has been historically problematic in comparative studies for several reasons, many pertaining to links between the proposed innovation and the formation of reproductive isolation between diverging lineages (Gillespie et al., 2020; Rabosky, 2017). Overall, this study agrees with the findings of Garcia-Porta and Ord (2013) that the presence of toepads is not a strong predictor of whether diplodactyloids colonize islands. This lack of relationship holds true for caudal scansors as well. The failure to find a relationships between diversification and adhesive scansors is contradictory to the traditional view that these structures have helped geckos and other padded lizards to colonize niches inaccessible to non-padded forms (Autumn & Peattie, 2002; Hansen & Autumn, 2005). The apparent absence of links between these distinct morphologies and biogeography and between them and speciation is not surprising given that both traits have evolved multiple times across geckos and that the serial homology of these structures complicates inferences of their unique nature in disparate lineages (Gamble, Greenbaum, Jackman, Russell & Bauer, 2012; Griffing, Gamble, Cohn & Sanger, 2022; Griffing, Sanger, Epperlein, Bauer, Cobos, Higham, Naylor & Gamble, 2021; Russell & Gamble, 2019). In contrast, diurnality and viviparity are entirely absent in mainland diplodactyloids, suggesting that biogeography has facilitated niche expansion and/or these traits have played a role in *in-situ* diversification. Diurnality has evolved multiple times in geckos across geographic settings, particularly amongst sphaerodactylids and gekkonids, though few studies have explicitly tested if it has acted as a key innovation in speciose lineages like sphearodactylines, *Cnemaspis*, and *Phelsuma*. Gamble, Greenbaum, Jackman and Bauer (2015) suggests that shifts in diel pattern may be driven by a myriad of factors including thermoregulation, predation, and competition, particularly in regions with climactic extremes and a paucity of other diurnal squamates. Diurnal representatives amongst the Tasmantis lineages are limited to two morphologically conservative genera and it is entirely feasible that this trait evolved so recently that these lineages have not had sufficient time to expand into additional diurnal niches.

## Conclusions

By leveraging ecomorphological data, natural history, biogeography, and abiotic niche evolution, we demonstrate that ecology and life history have driven diversification in diplodactyloid geckos. We highlighted axes upon which selection may be acting in driving phenotypic diversification before assessing if this diversity was driven by interspecific interactions or extrinsic abiotic factors. The two lineages of Zealandian diplodactyloid appear to have converged on similar phenotypes on multiple instances, that shifts in ecomorphology closely correspond to shifts in ecology, and that abiotic factors are a major component of lineage diversification in these adaptive radiations.

## Data Availability

The raw genomic data of this study are available on the Mendeley Data Platform (DOI:10.17632/zstmhjfkw5.1). Raw, untransformed meristic data used in morphometric analyses are available at (https://academic.oup.com/evolut/article/77/1/138/6881553#supplementary-data). Additional supporting can be found in the supplement for this paper on the publisher’s website.

## Acknowledgements and Funding

We would like to thank Jimmy McGuire, Paul Oliver, Sean Rovito, and Guin Wogan for their advice on structuring the early stages if this project. Special thanks to the collections staff at the Museum of Vertebrate Zoology, Western Australian Museum, Australian Museum, Queensland Museum, South Australian Museum, Museum of Victoria, Australia National University, Auckland Museum, and Te Papa Museum for providing tissues and granting us access to their respective collections. Funding awarded to PLS was provided by the National Science Foundation (DEB 1601806).

## Author Contributions

P.L.S. conceived of this project, collected all molecular and morphometric data, conducted most data analysis, and primarily wrote the manuscript. N.CR and R.ZF contributed to data analysis and were integral in manuscript edits.

## Supporting Information

Additional supporting information may be found on the publisher’s website.

**Skipwith_EJLS_Supp_mat**. Single file including additional tables and figures that contrast the results obtained through cPCA and pPCA on meristic data.

---

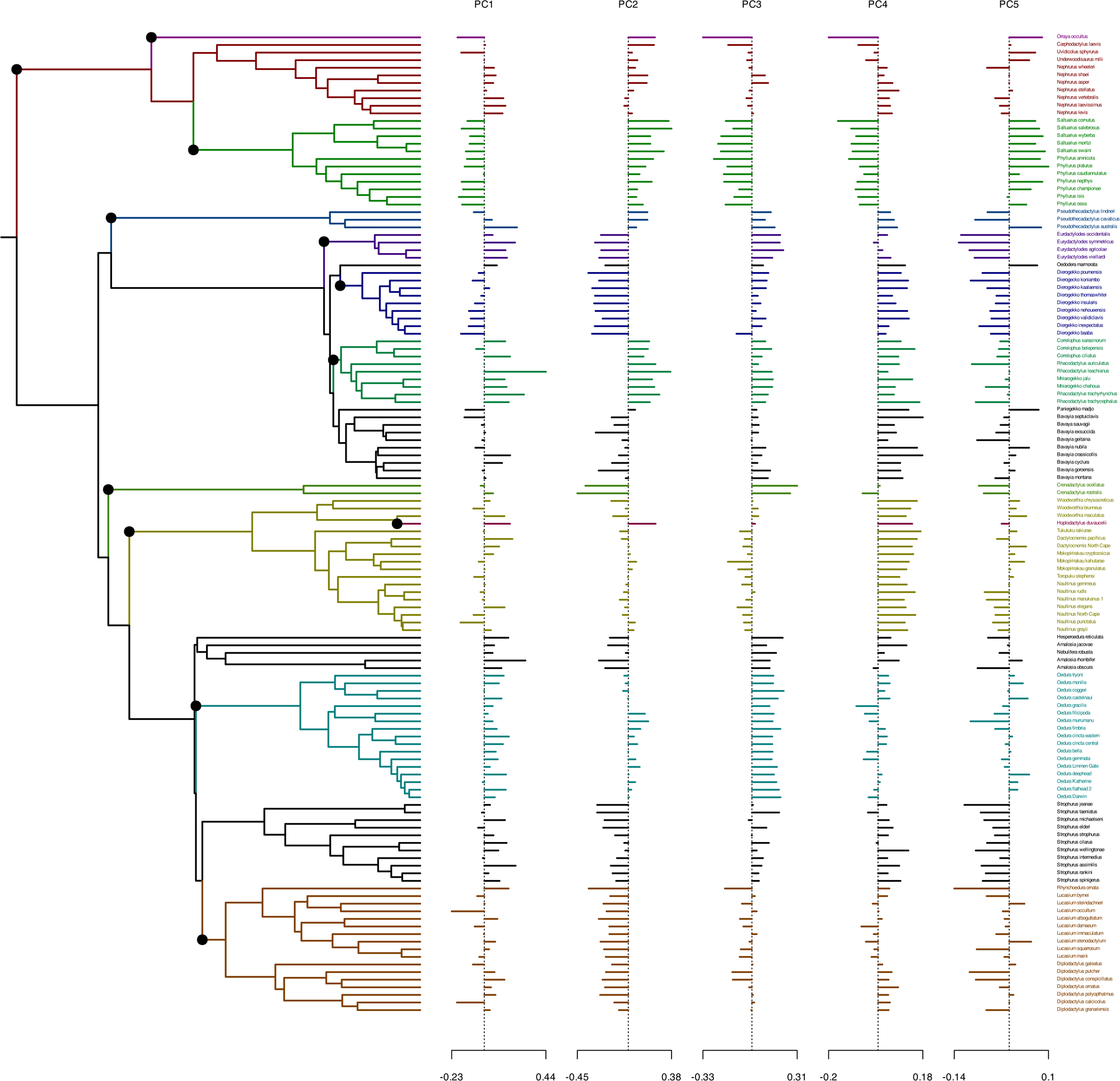

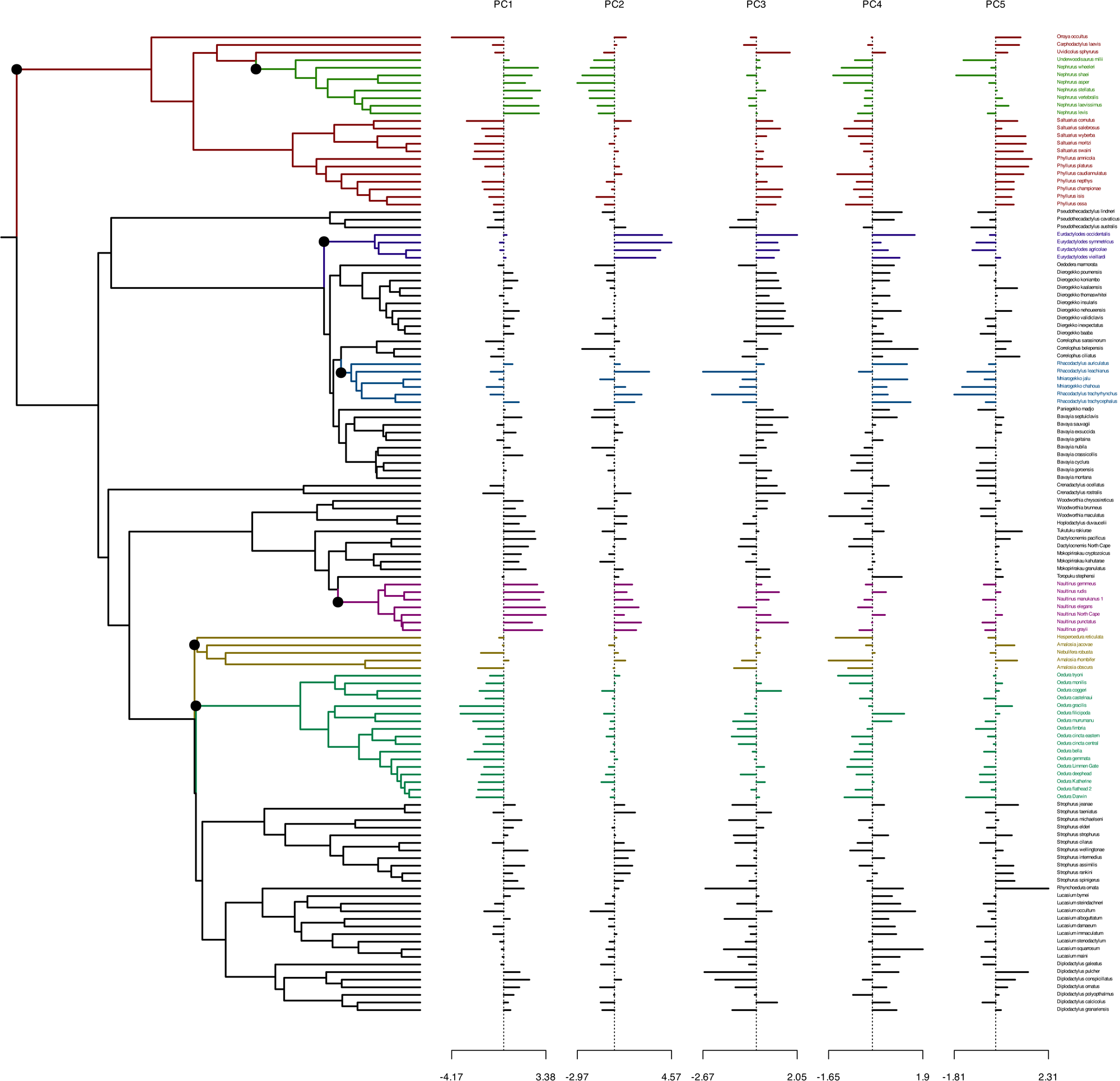

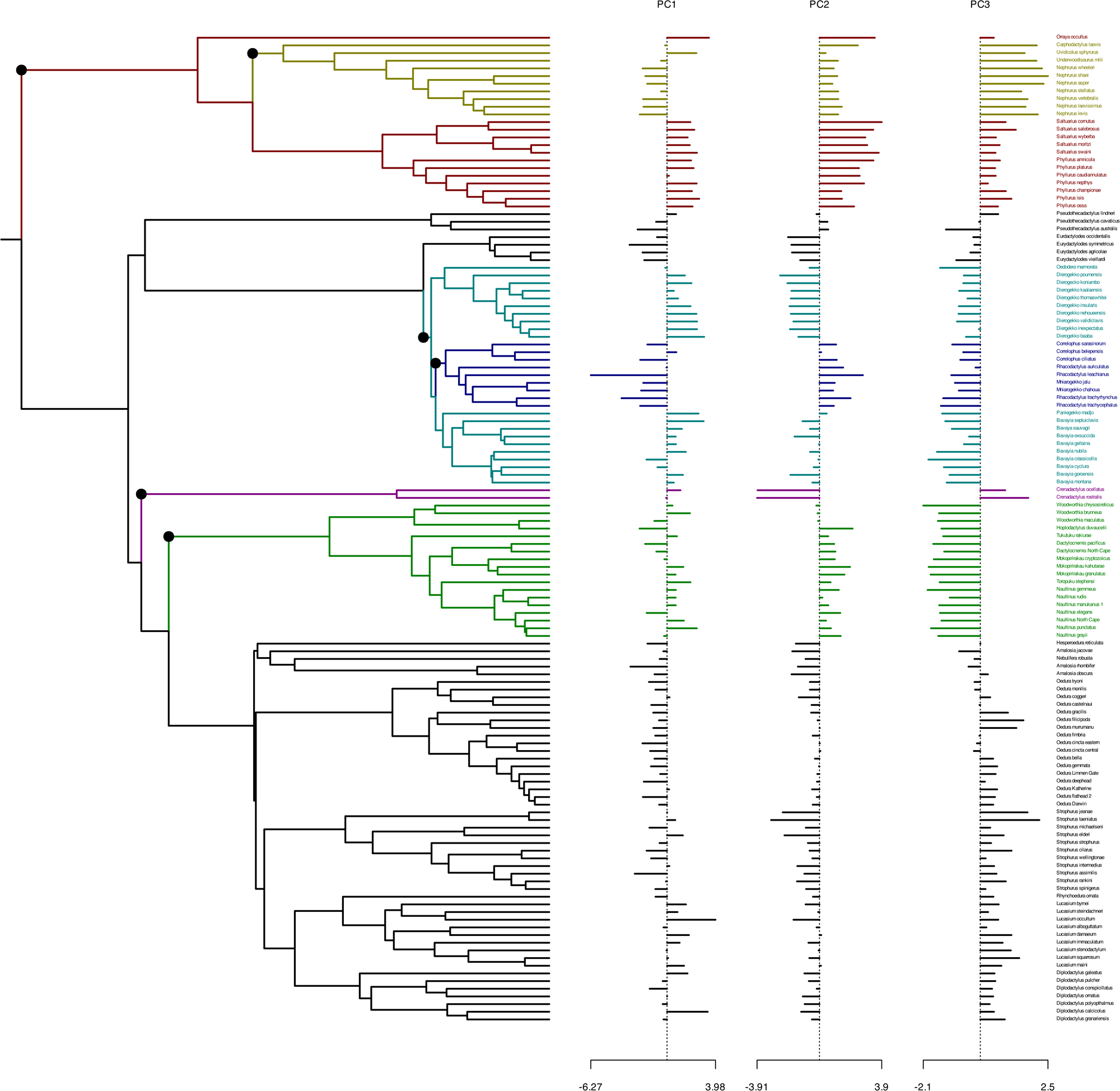

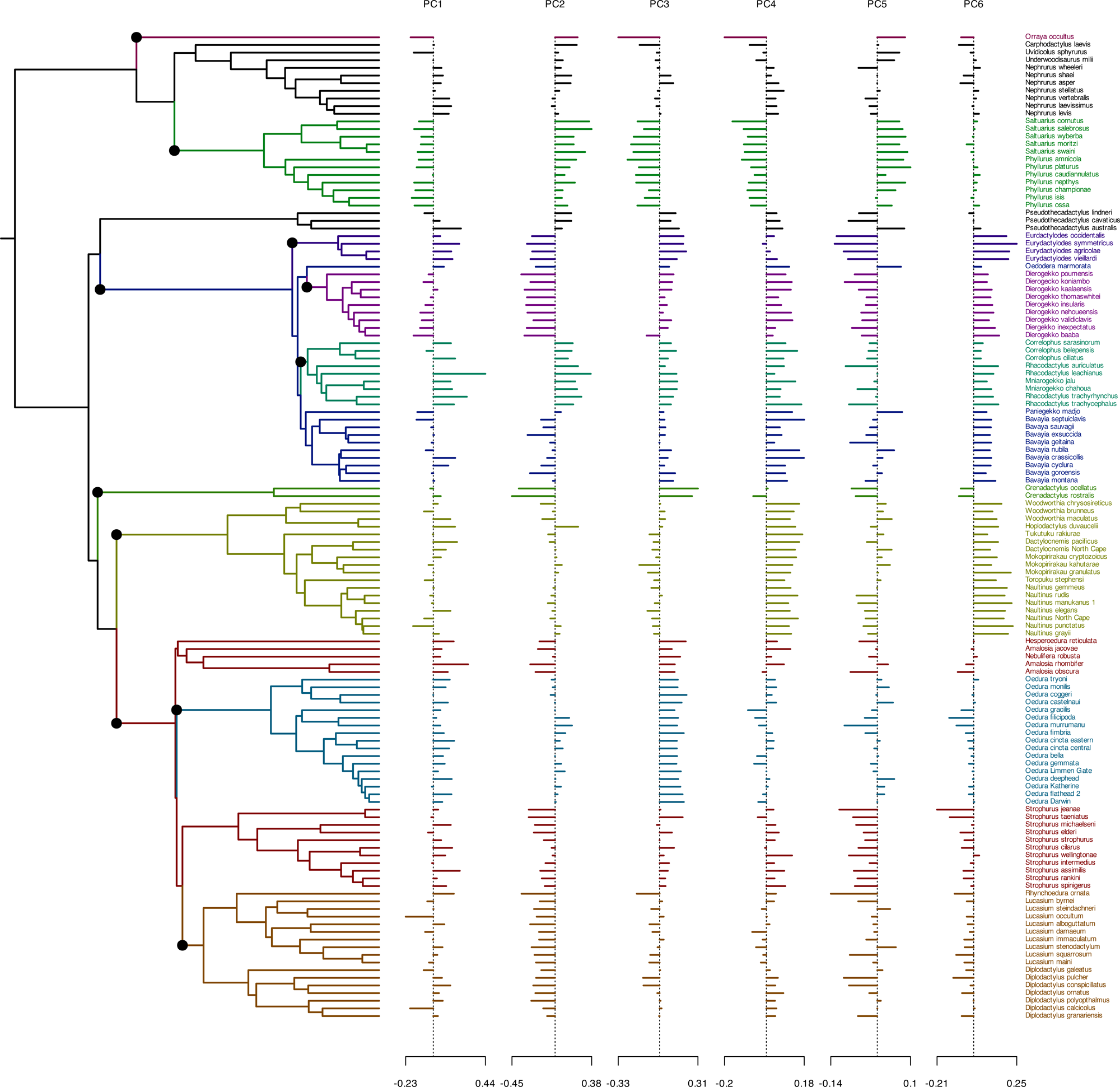

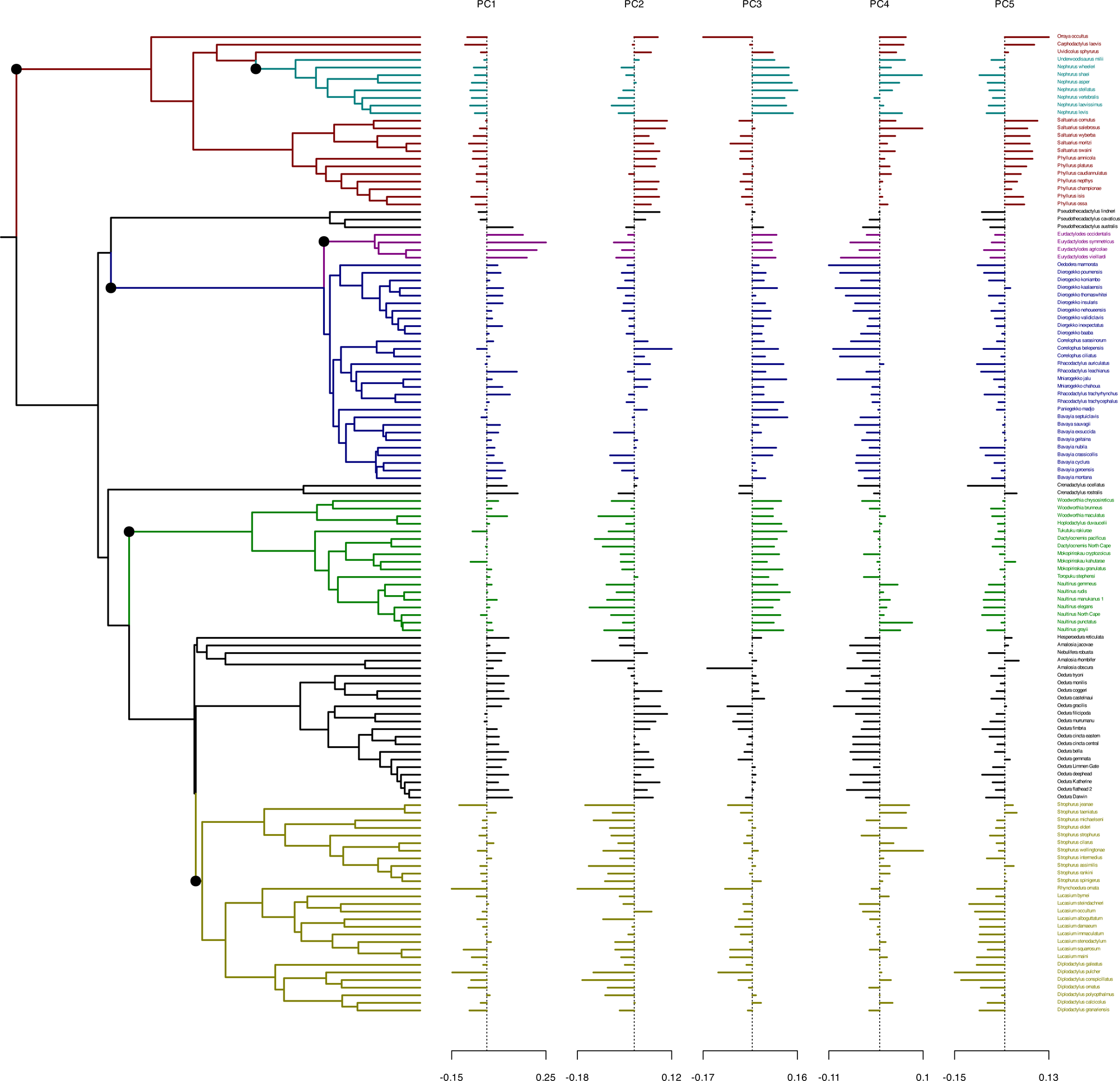

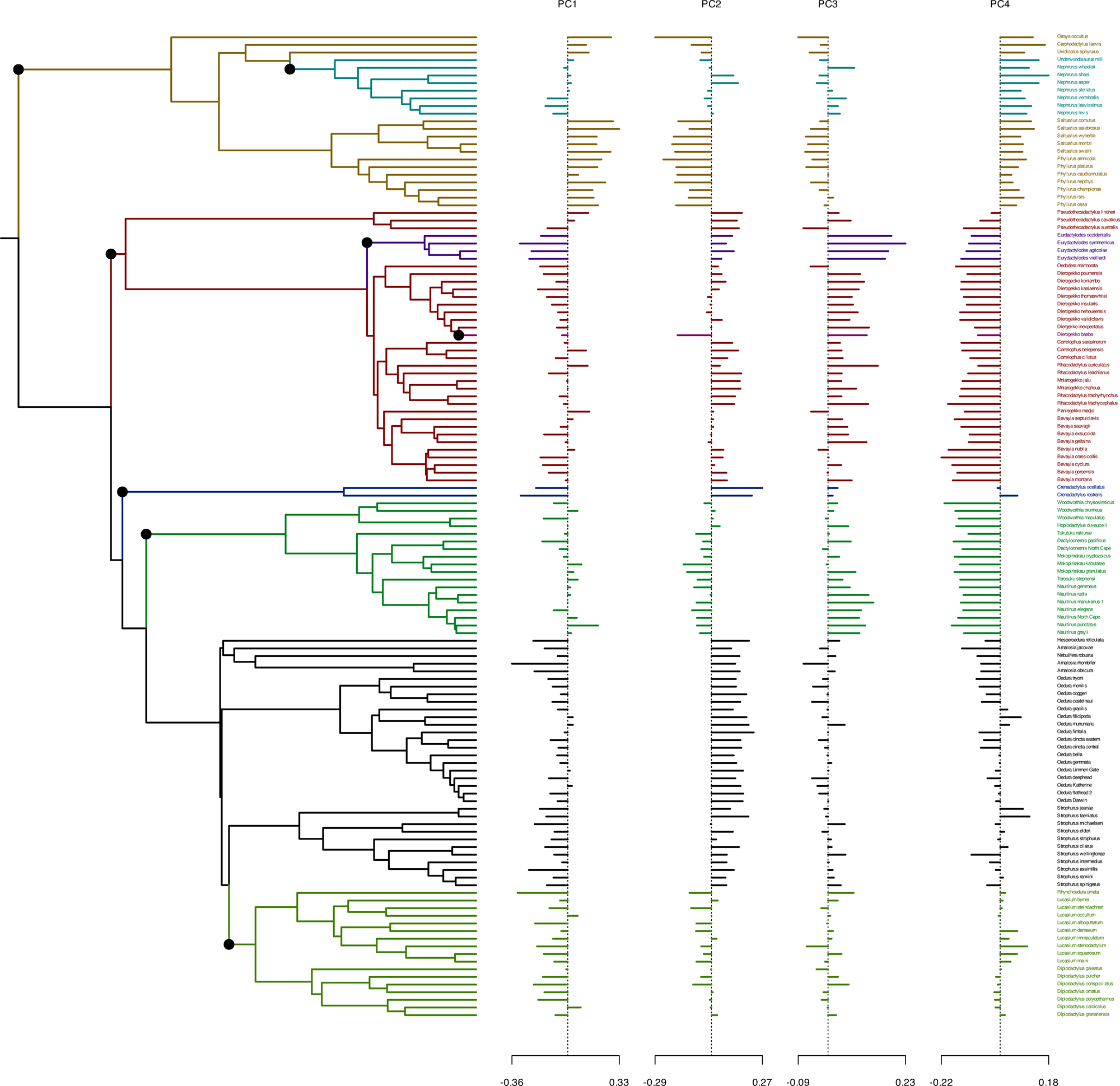

## Notes

DOI:10.17632/zstmhjfkw5.1

https://academic.oup.com/evolut/article/77/1/138/6881553#supplementary-data

### Competing Interest Statement

The authors have declared no competing interest.

