## Supplementary Material for "Abiotic Factors and Competitive Exclusion Drive Assembly Patterns in Two Insular Gecko Adaptive Radiations Displaying Ecomorphological Convergence"

### Methods:

**Supplementary Table S1**: Measurements used in this study.

| | |
|---|---|
| **SVL** | Snout-vent length |
| **HW** | head width at widest point |
| **HD** | head depth at widest point |
| **HL** | head length from snout tip to anterior edge of ear |
| **HL-ret** | head length from snout tip to retroarticular process |
| **Trk** | trunk length from armpit to groin |
| **Crus** | elbow to wrist length |
| **Tibia** | knee to ankle length |
| **EN** | internarial distance from anterior edge of nares |
| **NEYE** | distance from posterior nares edge to anterior corner of eye |
| **EYE** | eye diameter |
| **EyeEar** | posterior eye corner to anterior edge of aural canal |
| **IE** | distance between anterior eye corners |
| **EarRet** | posterior edge of aural canal to retroarticular process |
| **4FW** | length of 4th finger from claw to palm |
| **4FL** | width of 4th finger at widest point |
| **4TW** | length of 4th toe from claw to palm |
| **4TL** | width of 4th toe at widest point |

*PhyloPath Analyses*

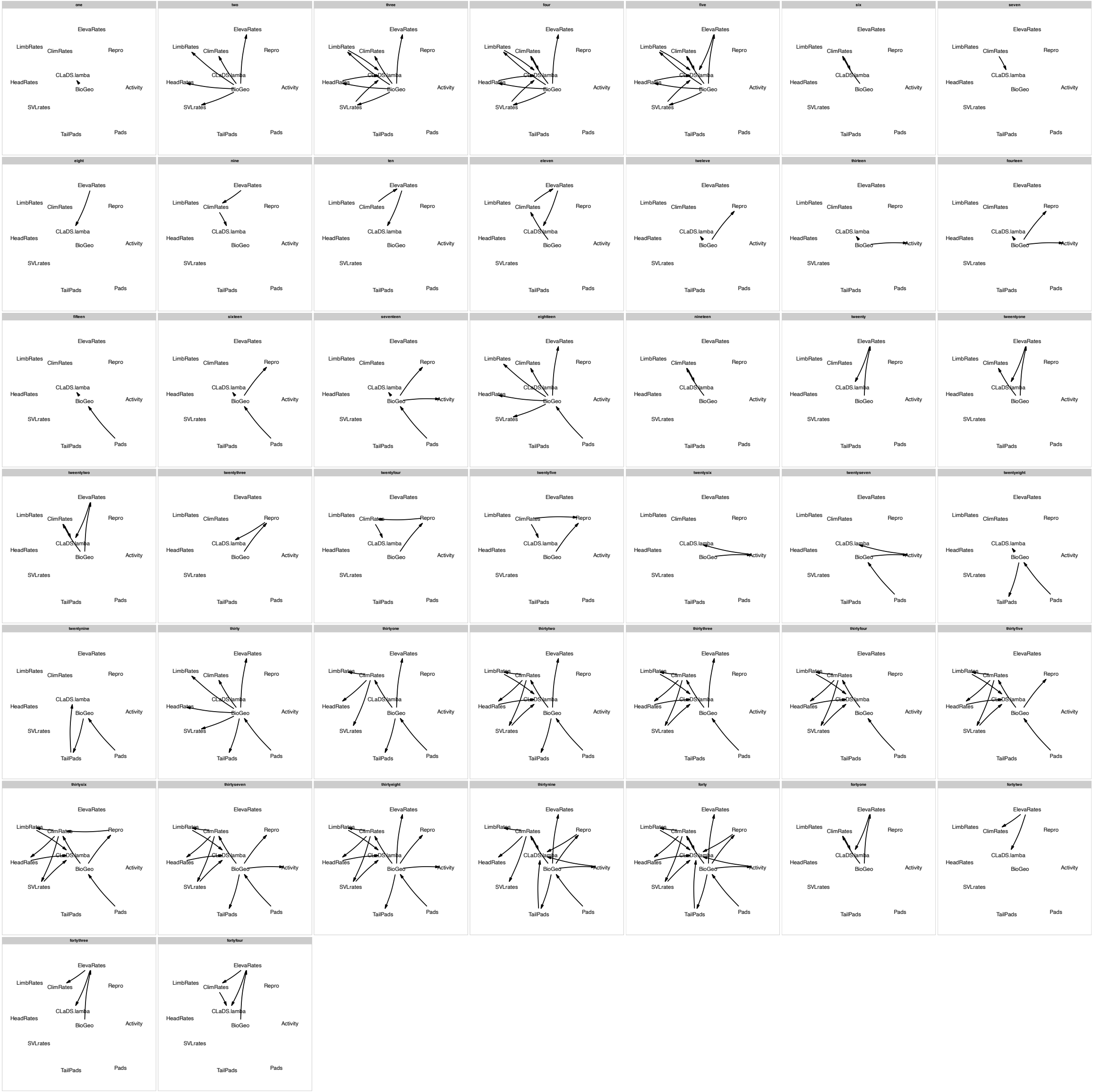


*Figure 1: Forty-four models used phylogenetic path analysis of the broader diplodactyloid clade (limbed forms only)*

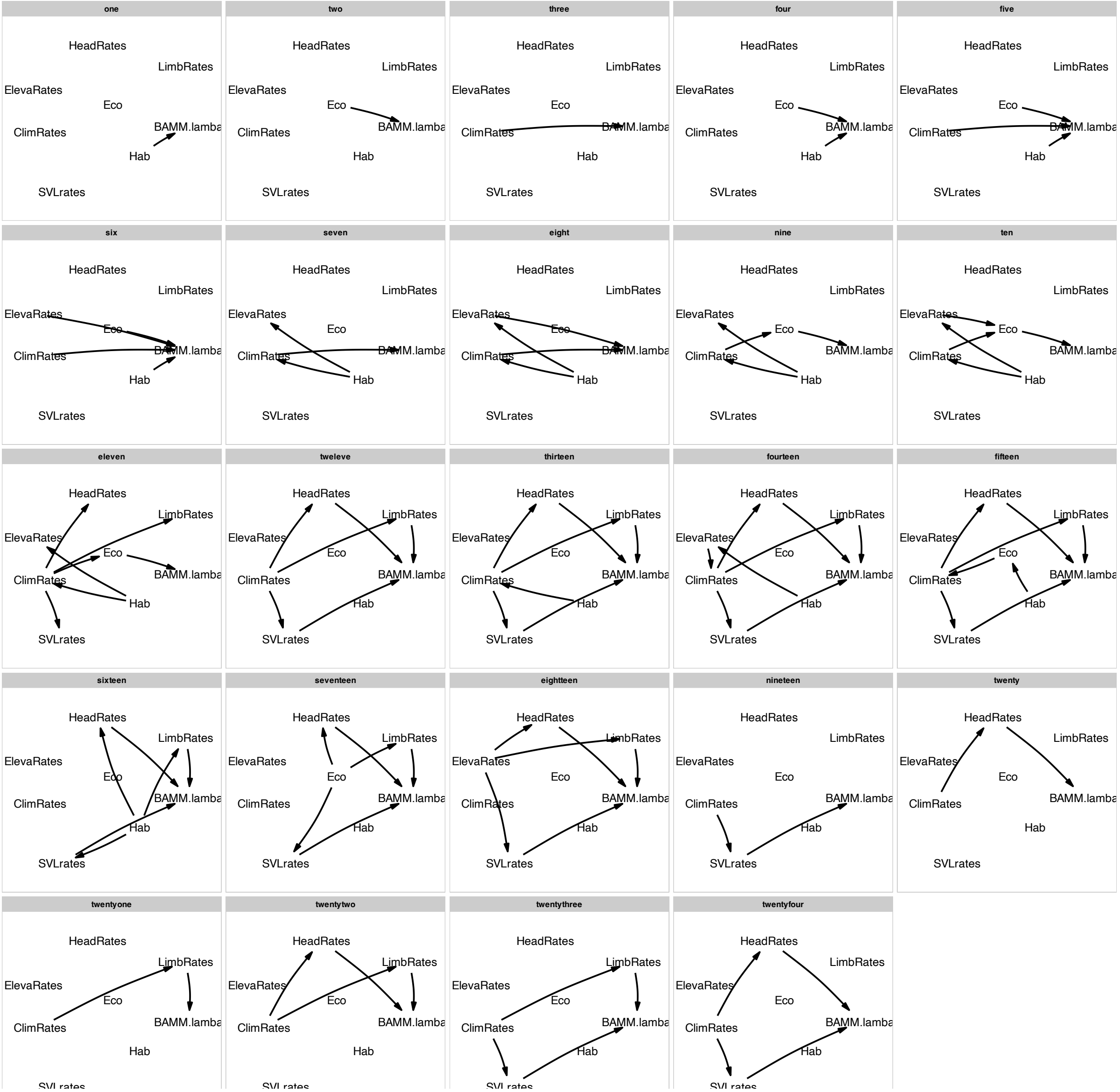


*Figure 2: Twenty-four models used in phylogenetic path analysis of the individual New Caledonian and New Zealand clades*

### Results:

**Supplementary Table S2:** Results of conventional principal component analyses (cPCA) and phylogenetic principal component analyses (pPCA) for all datasets.

| | Dataset | PC | Trait | Variance explained (%) |
|---|---|---|---|---|
| cPCA | Cranial | PC1 | HL.ret | 38.2 |
| | | PC2 | EYE.HL | 21.9 |
| | | PC3 | HD.HL | 13.3 |
| | | PC4 | HW.HL | 8.5 |
| | | PC5 | HD.HL | 7.5 |
| | Limbs | PC1 | FW.FL | 43.5 |
| | | PC2 | TW.TL | 33.1 |
| | | PC3 | Fing.Len | 18.3 |
| pPCA | Cranial | PC1 | EYE.HL | 28.1 |
| | | PC2 | HLret | 23.0 |
| | | PC3 | EYE.HL | 18.6 |
| | | PC4 | HD.HL | 12.9 |
| | | PC5 | EYE.HL | 11.2 |
| | Limbs | PC1 | Crus.SVL | 54.2 |
| | | PC2 | TW.TL | 21.6 |

| | | |
|---|---|---|
| PC3 | Fing.Len.SVL | 10.8 |
| PC4 | Tibia.SVL | 8.6 |
| PC5 | TW.TL | 3.7 |

**Supplementary Table S3**: Mean disparity indices (MDI) for both of the Zealandian clades with p-values from 1,000 simulations under BM. Bold p-values indicate significance. Estimates are for the date driven UCE tree only. Each clade was analyses individually. All estimates are for phylogenetic PCA (pPCA).

| | | New Caledonia | New Zealand |
|---|---|---|---|
| SVL | MDI | -0.26 | 1.25 |
| | p | **0.00** | 1.0 |
| Head only (cPCA-cranial: PCs 1-5) | MDI | 0.06 | 0.02 |
| | p | 0.75 | 0.52 |
| Limbed only (cPCA-limbed: PCs 1-3) | MDI | -0.12 | 0.04 |
| | p | > 0.05 | 0.56 |

**Supplementary Table S4**: Parameter estimates of RRphylo tests of convergence (search.conv). Contrasted are the cranial and limb proportion datasets for each the three *a-priori* ecological guilds. Of particular value are the parameters *ang.state.time* and *p.ang.state.time*, which are stronger indicators of the presence of convergent evolution. All estimates are for phylogenetic PCA (pPCA).

| Trait | State1 | State2 | ang.state | ang.state.time | p.ang.state | p.ang.state.time |
|---|---|---|---|---|---|---|
| cranial | NZ arboreal | NC arboreal | 44.94 | 0.65 | **0.00** | **0.02** |
| cranial | NZ scansorial | NC scansorial | 54.83 | 1.22 | **0.00** | 1.00 |
| cranial | NZ scansorial/rupicolous | NC scansorial/rupicolous | 52.54 | 0.78 | **0.01** | 0.37 |
| limbs | NZ arboreal | NC arboreal | 48.71 | 0.65 | **0.01** | 0.07 |
| limbs | NZ scansorial | NC scansorial | 52.83 | 1.23 | **0.00** | 1.00 |
| limbs | NZ scansorial/rupicolous | NC scansorial/rupicolous | 33.34 | 0.50 | **0.00** | **0.03** |

**Supplementary Table S5**: Parameter estimates of the Wheatsheaf index (*w*) from Windex. *w* is a metric for assessing the strength of convergence, not the presence of convergence (see results from RRphylo). Contrasted are the cranial and limb proportion datasets for each the three *a-priori* ecological guilds. All estimates are for phylogenetic PCA (pPCA).

| **Analysis** | **State** | ***w*** | **low95** | **up95** | **P** |
|---|---|---|---|---|---|
| all traits corrected for SVL | arboreal | 0.72 | 0.69 | 0.73 | 0.61 |
| all traits corrected for SVL | scansorial | 0.67 | 0.66 | 0.69 | 0.26 |
| all traits corrected for SVL | scansorial/rupicolous | 1.04 | 1.03 | 1.07 | **0.01** |
| cranial | arboreal | 1.08 | 1.06 | 1.12 | **0.00** |
| cranial | scansorial | 0.76 | 0.75 | 0.77 | **0.01** |
| cranial | scansorial/rupicolous | 1.29 | 1.27 | 1.34 | **0.00** |
| limbs | arboreal | 1.10 | 1.09 | 1.13 | **0.00** |
| limbs | scansorial | 0.75 | 0.73 | 0.77 | **0.02** |
| limbs | scansorial/rupicolous | 1.20 | 1.19 | 1.24 | **0.00** |

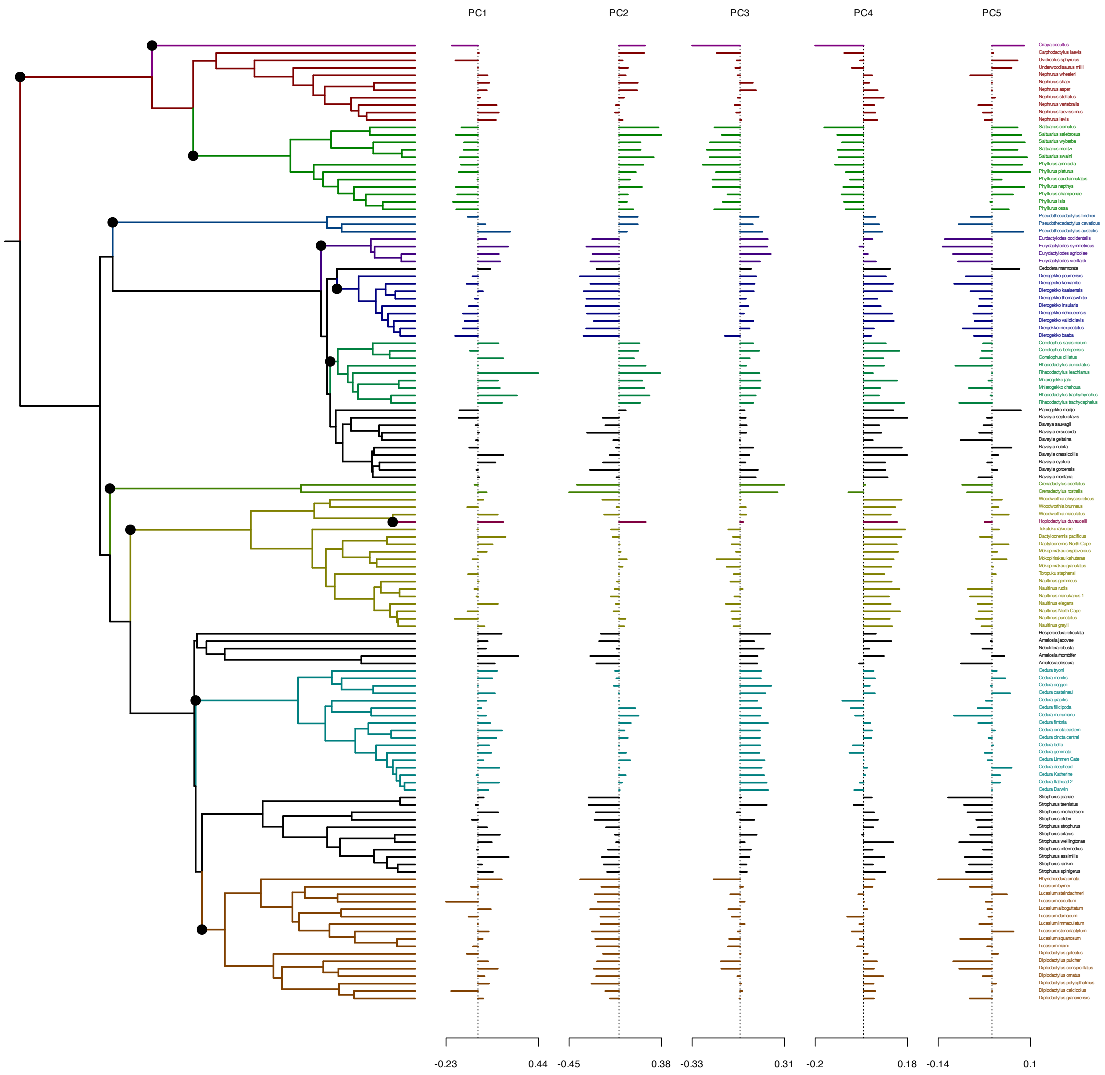

PC1
PC2
PC3
PC4
PC5
Orraya occultus
Carphodactylus laevis
Uvidicolus sphyrurus
Underwoodisaurus milii
Nephrurus wheeleri
Nephrurus shaei
Nephrurus asper
Nephrurus stellatus
Nephrurus vertebralis
Nephrurus laevissimus
Nephrurus levis
Saltuarius cornutus
Saltuarius salebrosus
Saltuarius wyberba
Saltuarius moritzi
Saltuarius swaini
Phyllurus amnicola
Phyllurus platurus
Phyllurus caudiannulatus
Phyllurus nepthys
Phyllurus championae
Phyllurus isis
Phyllurus ossa
Pseudothecadactylus lindneri
Pseudothecadactylus cavaticus
Pseudothecadactylus australis
Eurdactylodes occidentalis
Eurydactylodes symmetricus
Eurydactylodes agricolae
Eurydactylodes vieillardi
Oedodera marmorata
Dierogekko poumensis
Dierogecko koniambo
Dierogekko kaalaensis
Dierogekko thomaswhitei
Dierogekko insularis
Dierogekko nehoueensis
Dierogekko validiclavis
Diergekko inexpectatus
Dierogekko baaba
Correlophus sarasinorum
Correlophus belepensis
Correlophus ciliatus
Rhacodactylus auriculatus
Rhacodactylus leachianus
Mniarogekko jalu
Mniarogekko chahoua
Rhacodactylus trachyrhynchus
Rhacodactylus trachycephalus
Paniegekko madjo
Bavayia septuiclavis
Bavaya sauvagii
Bavayia exsuccida
Bavayia geitaina
Bavayia nubila
Bavayia crassicollis
Bavayia cyclura
Bavayia goroensis
Bavayia montana
Crenadactylus ocellatus
Crenadactylus rostralis
Woodworthia chrysosireticus
Woodworthia brunneus
Woodworthia maculatus
Hoplodactylus duvaucelii
Tukutuku rakiurae
Dactylocnemis pacificus
Dactylocnemis North Cape
Mokopirirakau cryptozoicus
Mokopirirakau kahutarae
Mokopirirakau granulatus
Toropuku stephensi
Naultinus gemmeus
Naultinus rudis
Naultinus manukanus 1
Naultinus elegans
Naultinus North Cape
Naultinus punctatus
Naultinus grayii
Hesperoedura reticulata
Amalosia jacovae
Nebulifera robusta
Amalosia rhombifer
Amalosia obscura
Oedura tryoni
Oedura monilis
Oedura coggeri
Oedura castelnaui
Oedura gracilis
Oedura filicipoda
Oedura murrumanu
Oedura fimbria
Oedura cincta eastern
Oedura cincta central
Oedura bella
Oedura gemmata
Oedura Limmen Gate
Oedura deephead
Oedura Katherine
Oedura flathead 2
Oedura Darwin
Strophurus jeanae
Strophurus taeniatus
Strophurus michaelseni
Strophurus elderi
Strophurus strophurus
Strophurus cilarus
Strophurus wellingtonae
Strophurus intermedius
Strophurus assimilis
Strophurus rankini
Strophurus spinigerus
Rhynchoedura ornata
Lucasium byrnei
Lucasium steindachneri
Lucasium occultum
Lucasium alboguttatum
Lucasium damaeum
Lucasium immaculatum
Lucasium stenodactylum
Lucasium squarrosum
Lucasium maini
Diplodactylus galeatus
Diplodactylus pulcher
Diplodactylus conspicillatus
Diplodactylus ornatus
Diplodactylus polyopthalmus
Diplodactylus calcicolus
Diplodactylus granariensis
-0.23
0.44
-0.45
0.38
-0.33
0.31
-0.2
0.18
-0.14
0.1

*Figure 3: Phylogenetic EM analysis of all traits corrected for SVL using cPCA. Node circles indicate where adaptive shifts were identified.*

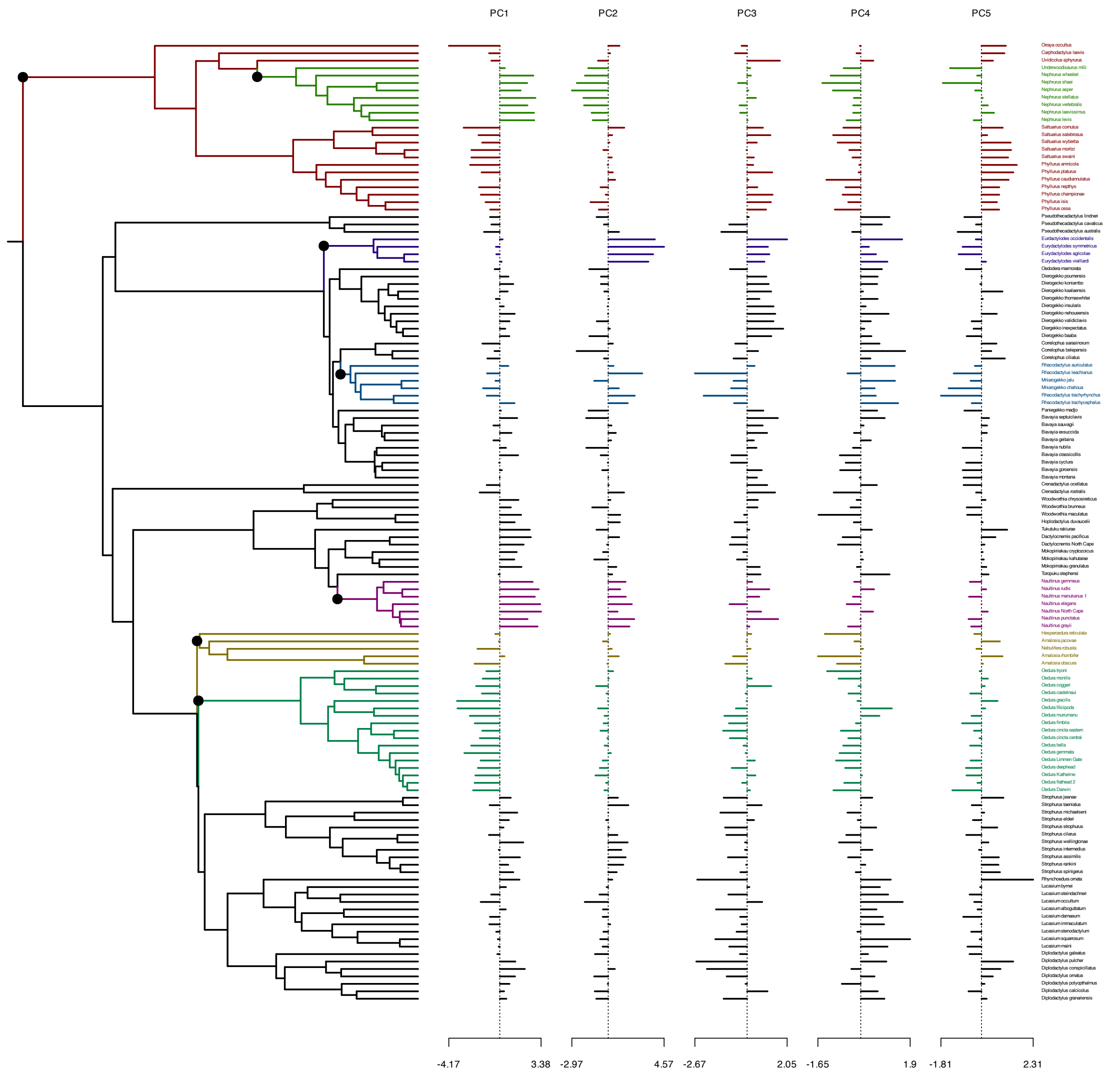
PC1
PC2
PC3
PC4
PC5
Orraya occultus
Carphodactylus laevis
Uvidicolus sphyrurus
Underwoodisaurus milii
Nephrurus wheeleri
Nephrurus shaei
Nephrurus asper
Nephrurus stellatus
Nephrurus vertebralis
Nephrurus laevissimus
Nephrurus levis
Saltuarius cornutus
Saltuarius salebrosus
Saltuarius wyberba
Saltuarius moritzi
Saltuarius swaini
Phyllurus amnicola
Phyllurus platurus
Phyllurus caudiannulatus
Phyllurus nepthys
Phyllurus championae
Phyllurus isis
Phyllurus ossa
Pseudothecadactylus lindneri
Pseudothecadactylus cavaticus
Pseudothecadactylus australis
Eurdactylodes occidentalis
Eurydactylodes symmetricus
Eurydactylodes agricolae
Eurydactylodes vieillardi
Oedodera marmorata
Dierogekko poumensis
Dierogecko koniambo
Dierogekko kaalaensis
Dierogekko thomaswhitei
Dierogekko insularis
Dierogekko nehoueensis
Dierogekko validiclavis
Diergekko inexpectatus
Dierogekko baaba
Correlophus sarasinorum
Correlophus belepensis
Correlophus ciliatus
Rhacodactylus auriculatus
Rhacodactylus leachianus
Mniarogekko jalu
Mniarogekko chahoua
Rhacodactylus trachyrhynchus
Rhacodactylus trachycephalus
Paniegekko madjo
Bavayia septuiclavis
Bavaya sauvagii
Bavayia exsuccida
Bavayia geitaina
Bavayia nubila
Bavayia crassicollis
Bavayia cyclura
Bavayia goroensis
Bavayia montana
Crenadactylus ocellatus
Crenadactylus rostralis
Woodworthia chrysosireticus
Woodworthia brunneus
Woodworthia maculatus
Hoplodactylus duvaucelii
Tukutuku rakiurae
Dactylocnemis pacificus
Dactylocnemis North Cape
Mokopirirakau cryptozoicus
Mokopirirakau kahutarae
Mokopirirakau granulatus
Toropuku stephensi
Naultinus gemmeus
Naultinus rudis
Naultinus manukanus 1
Naultinus elegans
Naultinus North Cape
Naultinus punctatus
Naultinus grayii
Hesperoedura reticulata
Amalosia jacovae
Nebulifera robusta
Amalosia rhombifer
Amalosia obscura
Oedura tryoni
Oedura monilis
Oedura coggeri
Oedura castelnaui
Oedura gracilis
Oedura filicipoda
Oedura murrumanu
Oedura fimbria
Oedura cincta eastern
Oedura cincta central
Oedura bella
Oedura gemmata
Oedura Limmen Gate
Oedura deephead
Oedura Katherine
Oedura flathead 2
Oedura Darwin
Strophurus jeanae
Strophurus taeniatus
Strophurus michaelseni
Strophurus elderi
Strophurus strophurus
Strophurus cilarus
Strophurus wellingtonae
Strophurus intermedius
Strophurus assimilis
Strophurus rankini
Strophurus spinigerus
Rhynchoedura ornata
Lucasium byrnei
Lucasium steindachneri
Lucasium occultum
Lucasium alboguttatum
Lucasium damaeum
Lucasium immaculatum
Lucasium stenodactylum
Lucasium squarrosum
Lucasium maini
Diplodactylus galeatus
Diplodactylus pulcher
Diplodactylus conspicillatus
Diplodactylus ornatus
Diplodactylus polyopthalmus
Diplodactylus calcicolus
Diplodactylus granariensis
-4.17
3.38
-2.97
4.57
-2.67
2.05
-1.65
1.9
-1.81
2.31

*Figure 4: Phylogenetic EM analysis of cranial traits using cPCA. Node circles indicate where adaptive shifts were identified.*

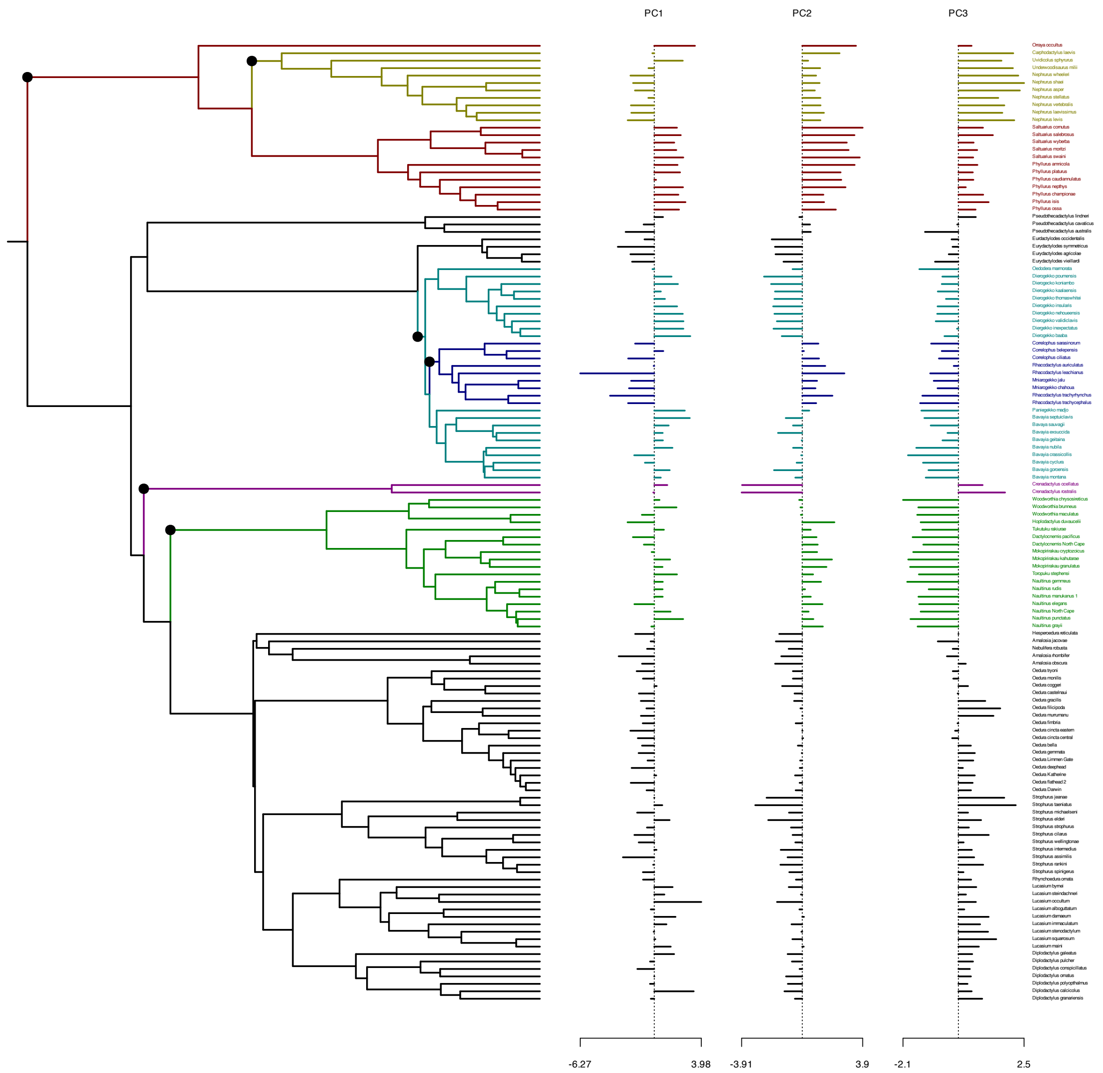
PC1
PC2
PC3
Orraya occultus
Carphodactylus laevis
Uvidicolus sphyrurus
Underwoodisaurus milii
Nephrurus wheeleri
Nephrurus shaei
Nephrurus asper
Nephrurus stellatus
Nephrurus vertebralis
Nephrurus laevissimus
Nephrurus levis
Saltuarius cornutus
Saltuarius salebrosus
Saltuarius wyberba
Saltuarius moritzi
Saltuarius swaini
Phyllurus amnicola
Phyllurus platurus
Phyllurus caudiannulatus
Phyllurus nepthys
Phyllurus championae
Phyllurus isis
Phyllurus ossa
Pseudothecadactylus lindneri
Pseudothecadactylus cavaticus
Pseudothecadactylus australis
Eurdactylodes occidentalis
Eurydactylodes symmetricus
Eurydactylodes agricolae
Eurydactylodes vieillardi
Oedodera marmorata
Dierogekko poumensis
Dierogecko koniambo
Dierogekko kaalaensis
Dierogekko thomaswhitei
Dierogekko insularis
Dierogekko nehoueensis
Dierogekko validiclavis
Diergekko inexpectatus
Dierogekko baaba
Correlophus sarasinorum
Correlophus belepensis
Correlophus ciliatus
Rhacodactylus auriculatus
Rhacodactylus leachianus
Mniarogekko jalu
Mniarogekko chahoua
Rhacodactylus trachyrhynchus
Rhacodactylus trachycephalus
Paniegekko madjo
Bavayia septuiclavis
Bavaya sauvagii
Bavayia exsuccida
Bavayia geitaina
Bavayia nubila
Bavayia crassicollis
Bavayia cyclura
Bavayia goroensis
Bavayia montana
Crenadactylus ocellatus
Crenadactylus rostralis
Woodworthia chrysosireticus
Woodworthia brunneus
Woodworthia maculatus
Hoplodactylus duvaucelii
Tukutuku rakiurae
Dactylocnemis pacificus
Dactylocnemis North Cape
Mokopirirakau cryptozoicus
Mokopirirakau kahutarae
Mokopirirakau granulatus
Toropuku stephensi
Naultinus gemmeus
Naultinus rudis
Naultinus manukanus 1
Naultinus elegans
Naultinus North Cape
Naultinus punctatus
Naultinus grayii
Hesperoedura reticulata
Amalosia jacovae
Nebulifera robusta
Amalosia rhombifer
Amalosia obscura
Oedura tryoni
Oedura monilis
Oedura coggeri
Oedura castelnaui
Oedura gracilis
Oedura filicipoda
Oedura murrumanu
Oedura fimbria
Oedura cincta eastern
Oedura cincta central
Oedura bella
Oedura gemmata
Oedura Limmen Gate
Oedura deephead
Oedura Katherine
Oedura flathead 2
Oedura Darwin
Strophurus jeanae
Strophurus taeniatus
Strophurus michaelseni
Strophurus elderi
Strophurus strophurus
Strophurus cilarus
Strophurus wellingtonae
Strophurus intermedius
Strophurus assimilis
Strophurus rankini
Strophurus spinigerus
Rhynchoedura ornata
Lucasium byrnei
Lucasium steindachneri
Lucasium occultum
Lucasium alboguttatum
Lucasium damaeum
Lucasium immaculatum
Lucasium stenodactylum
Lucasium squarrosum
Lucasium maini
Diplodactylus galeatus
Diplodactylus pulcher
Diplodactylus conspicillatus
Diplodactylus ornatus
Diplodactylus polyopthalmus
Diplodactylus calcicolus
Diplodactylus granariensis
-6.27
3.98
-3.91
3.9
-2.1
2.5

*Figure 5: Phylogenetic EM analysis of limb traits using cPCA. Node circles indicate where adaptive shifts were identified.*

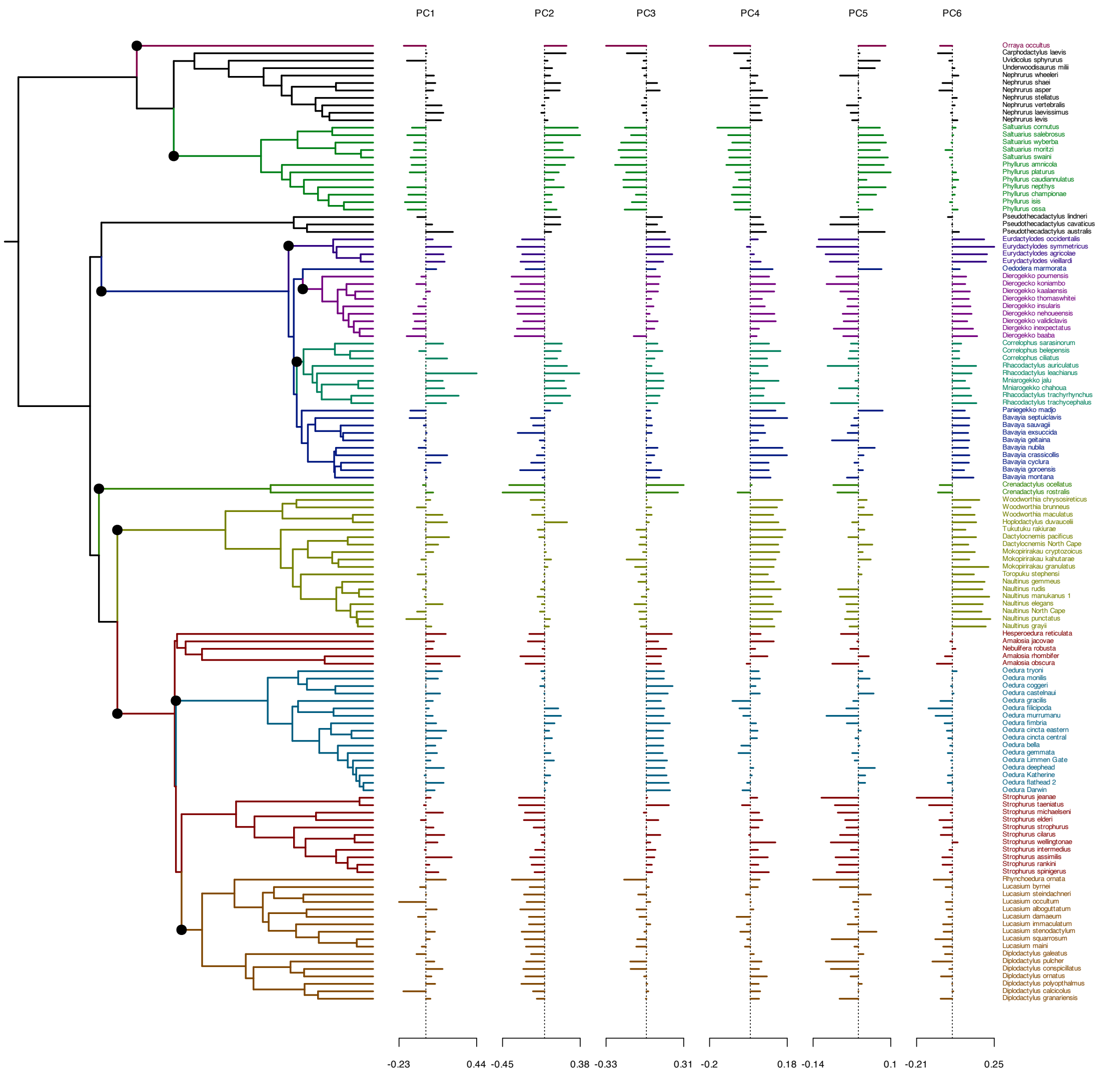
PC1
PC2
PC3
PC4
PC5
PC6
Orraya occultus
Carphodactylus laevis
Uvidicolus sphyrurus
Underwoodisaurus milii
Nephrurus wheeleri
Nephrurus shaei
Nephrurus asper
Nephrurus stellatus
Nephrurus vertebralis
Nephrurus laevissimus
Nephrurus levis
Saltuarius cornutus
Saltuarius salebrosus
Saltuarius wyberba
Saltuarius moritzi
Saltuarius swaini
Phyllurus amnicola
Phyllurus platurus
Phyllurus caudiannulatus
Phyllurus nepthys
Phyllurus championae
Phyllurus isis
Phyllurus ossa
Pseudothecadactylus lindneri
Pseudothecadactylus cavaticus
Pseudothecadactylus australis
Eurdactylodes occidentalis
Eurydactylodes symmetricus
Eurydactylodes agricolae
Eurydactylodes vieillardi
Oedodera marmorata
Dierogekko poumensis
Dierogecko koniambo
Dierogekko kaalaensis
Dierogekko thomaswhitei
Dierogekko insularis
Dierogekko nehoueensis
Dierogekko validiclavis
Diergekko inexpectatus
Dierogekko baaba
Correlophus sarasinorum
Correlophus belepensis
Correlophus ciliatus
Rhacodactylus auriculatus
Rhacodactylus leachianus
Mniarogekko jalu
Mniarogekko chahoua
Rhacodactylus trachyrhynchus
Rhacodactylus trachycephalus
Paniegekko madjo
Bavayia septuiclavis
Bavaya sauvagii
Bavayia exsuccida
Bavayia geitaina
Bavayia nubila
Bavayia crassicollis
Bavayia cyclura
Bavayia goroensis
Bavayia montana
Crenadactylus ocellatus
Crenadactylus rostralis
Woodworthia chrysosireticus
Woodworthia brunneus
Woodworthia maculatus
Hoplodactylus duvaucelii
Tukutuku rakiurae
Dactylocnemis pacificus
Dactylocnemis North Cape
Mokopirirakau cryptozoicus
Mokopirirakau kahutarae
Mokopirirakau granulatus
Toropuku stephensi
Naultinus gemmeus
Naultinus rudis
Naultinus manukanus 1
Naultinus elegans
Naultinus North Cape
Naultinus punctatus
Naultinus grayii
Hesperoedura reticulata
Amalosia jacovae
Nebulifera robusta
Amalosia rhombifer
Amalosia obscura
Oedura tryoni
Oedura monilis
Oedura coggeri
Oedura castelnaui
Oedura gracilis
Oedura filicipoda
Oedura murrumanu
Oedura fimbria
Oedura cincta eastern
Oedura cincta central
Oedura bella
Oedura gemmata
Oedura Limmen Gate
Oedura deephead
Oedura Katherine
Oedura flathead 2
Oedura Darwin
Strophurus jeanae
Strophurus taeniatus
Strophurus michaelseni
Strophurus elderi
Strophurus strophurus
Strophurus cilarus
Strophurus wellingtonae
Strophurus intermedius
Strophurus assimilis
Strophurus rankini
Strophurus spinigerus
Rhynchoedura ornata
Lucasium byrnei
Lucasium steindachneri
Lucasium occultum
Lucasium alboguttatum
Lucasium damaeum
Lucasium immaculatum
Lucasium stenodactylum
Lucasium squarrosum
Lucasium maini
Diplodactylus galeatus
Diplodactylus pulcher
Diplodactylus conspicillatus
Diplodactylus ornatus
Diplodactylus polyopthalmus
Diplodactylus calcicolus
Diplodactylus granariensis
-0.23
0.44
-0.45
0.38
-0.33
0.31
-0.2
0.18
-0.14
0.1
-0.21
0.25

*Figure 6: Phylogenetic EM analysis of all traits corrected for SVL using pPCA. Node cirles indicate where adaptive shifts were identified.*

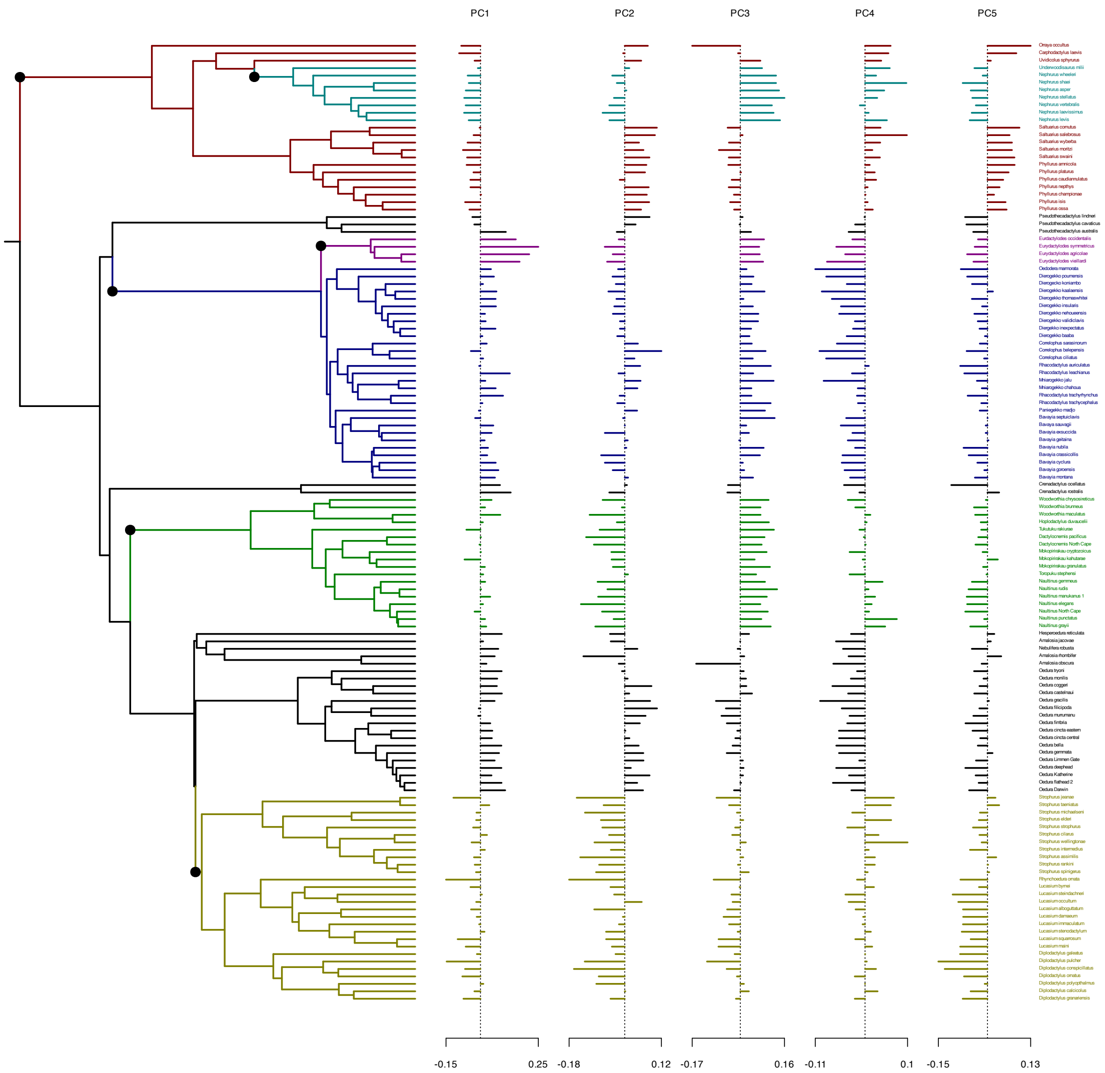
PC1
PC2
PC3
PC4
PC5
-0.15
0.25
-0.18
0.12
-0.17
0.16
-0.11
0.1
-0.15
0.13

*Figure 7: Phylogenetic EM analysis of cranial traits using pPCA. Node circles indicate where adaptive shifts were identified.*

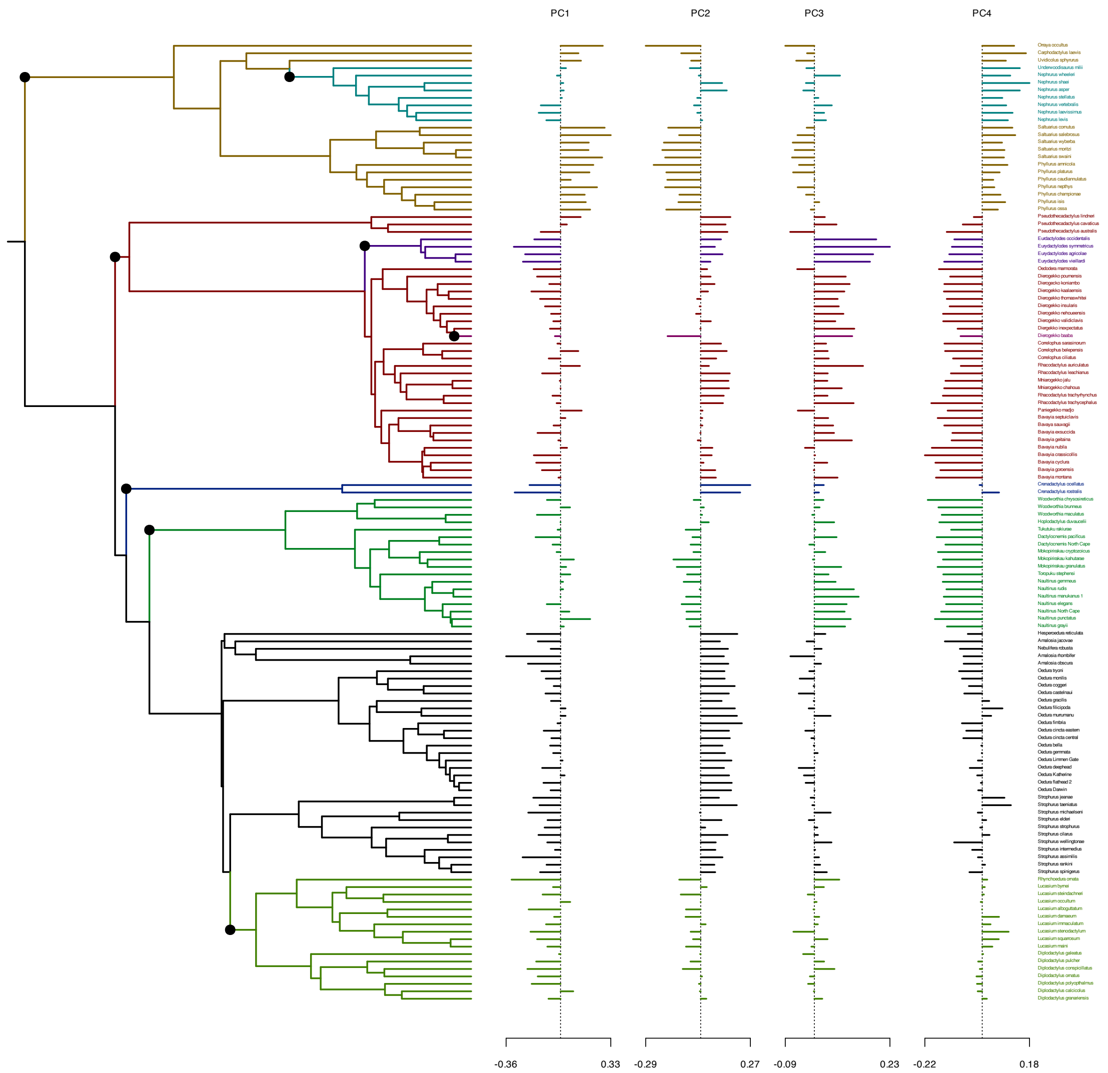

PC1
PC2
PC3
PC4
-0.36
0.33
-0.29
0.27
-0.09
0.23
-0.22
0.18
Orraya occultus
Carphodactylus laevis
Uvidicolus sphyrurus
Underwoodisaurus milii
Nephrurus wheeleri
Nephrurus shaei
Nephrurus asper
Nephrurus stellatus
Nephrurus vertebralis
Nephrurus laevissimus
Nephrurus levis
Saltuarius cornutus
Saltuarius salebrosus
Saltuarius wyberba
Saltuarius moritzi
Saltuarius swaini
Phyllurus amnicola
Phyllurus platurus
Phyllurus caudiannulatus
Phyllurus nepthys
Phyllurus championae
Phyllurus isis
Phyllurus ossa
Pseudothecadactylus lindneri
Pseudothecadactylus cavaticus
Pseudothecadactylus australis
Eurdactylodes occidentalis
Eurydactylodes symmetricus
Eurydactylodes agricolae
Eurydactylodes vieillardi
Oedodera marmorata
Dierogekko poumensis
Dierogecko koniambo
Dierogekko kaalaensis
Dierogekko thomaswhitei
Dierogekko insularis
Dierogekko nehoueensis
Dierogekko validiclavis
Diergekko inexpectatus
Dierogekko baaba
Correlophus sarasinorum
Correlophus belepensis
Correlophus ciliatus
Rhacodactylus auriculatus
Rhacodactylus leachianus
Mniarogekko jalu
Mniarogekko chahoua
Rhacodactylus trachyrhynchus
Rhacodactylus trachycephalus
Paniegekko madjo
Bavayia septuiclavis
Bavaya sauvagii
Bavayia exsuccida
Bavayia geitaina
Bavayia nubila
Bavayia crassicollis
Bavayia cyclura
Bavayia goroensis
Bavayia montana
Crenadactylus ocellatus
Crenadactylus rostralis
Woodworthia chrysosireticus
Woodworthia brunneus
Woodworthia maculatus
Hoplodactylus duvaucelii
Tukutuku rakiurae
Dactylocnemis pacificus
Dactylocnemis North Cape
Mokopirirakau cryptozoicus
Mokopirirakau kahutarae
Mokopirirakau granulatus
Toropuku stephensi
Naultinus gemmeus
Naultinus rudis
Naultinus manukanus 1
Naultinus elegans
Naultinus North Cape
Naultinus punctatus
Naultinus grayii
Hesperoedura reticulata
Amalosia jacovae
Nebulifera robusta
Amalosia rhombifer
Amalosia obscura
Oedura tryoni
Oedura monilis
Oedura coggeri
Oedura castelnaui
Oedura gracilis
Oedura filicipoda
Oedura murrumanu
Oedura fimbria
Oedura cincta eastern
Oedura cincta central
Oedura bella
Oedura gemmata
Oedura Limmen Gate
Oedura deephead
Oedura Katherine
Oedura flathead 2
Oedura Darwin
Strophurus jeanae
Strophurus taeniatus
Strophurus michaelseni
Strophurus elderi
Strophurus strophurus
Strophurus cilarus
Strophurus wellingtonae
Strophurus intermedius
Strophurus assimilis
Strophurus rankini
Strophurus spinigerus
Rhynchoedura ornata
Lucasium byrnei
Lucasium steindachneri
Lucasium occultum
Lucasium alboguttatum
Lucasium damaeum
Lucasium immaculatum
Lucasium stenodactylum
Lucasium squarrosum
Lucasium maini
Diplodactylus galeatus
Diplodactylus pulcher
Diplodactylus conspicillatus
Diplodactylus ornatus
Diplodactylus polyopthalmus
Diplodactylus calcicolus
Diplodactylus granariensis

*Figure 8: Phylogenetic EM analysis of limb traits using pPCA. Node circles indicate where adaptive shifts were identified.*
